# Generative AI-Guided Design of High-Affinity T Cell Receptors

**DOI:** 10.64898/2026.02.12.705547

**Authors:** Martin Renqiang Min, Tianxiao Li, Kazuhide Onoguchi, Daiki Mori, Ayako Demachi-Okamura, Jonathan Warrell, Pierre Machart, Anja Moesch, Andrea Meiser, Ivy Grace Pait, Daisuke Muraoka, Hirokazu Matsushita, Giulia Paiardi, Matheus Ferraz, Kaidre Bendjama

**Affiliations:** NEC Laboratories America, Princeton, New Jersey, United States; NEC Corporation, Tokyo, Japan; NEC Laboratories Europe GmbH, Heidelberg, Germany; Division of Translational Oncoimmunology, Aichi Cancer Center Research Institute, Nagoya, Japan; Division of Immune Response, Aichi Cancer Center Research Institute, Nagoya, Japan; NEC OncoImmunity AS, Oslo, Norway; NEC Bio, Schiphol, The Netherlands

**Author notes:** These authors contributed equally to this work (Martin Renqiang Min, Tianxiao Li, Kazuhide Onoguchi, Daiki Mori, Ayako Demachi-Okamura). Contributing authors.

**Keywords:** T cell receptor engineering, tumor antigen recognition, AI-driven protein design

## Abstract

Developing T cell receptors (TCRs) with sufficiently high affinity for tumor antigens (TAs) remains a fundamental challenge in TCR-T immunotherapy. Experimental methods such as affinity maturation and high-throughput screening have enabled the identification of TCRs with enhanced activity. However, their efficiency is often constrained by limited throughput, insufficient coverage, and the generally lower affinities of naturally occurring TCRs toward TAs. To address these challenges, we present `TCRPPO2`, an integrated AI-driven, *in silico* affinity maturation framework for peptide-specific TCR optimization. Using reinforcement learning, `TCRPPO2` learns mutation policies that iteratively enhance the TCR binding affinity to the target peptide, derived from predictive models trained on carefully curated interaction data. The model is further augmented by a generative AI critic model that discourages implausible designs to ensure the biophysical validity. The designs are further screened by robust post-screening methods that leverage diverse functional annotations and physical prior knowledge. We applied `TCRPPO2` to the clinically relevant MART-1 antigen and experimentally validated the designed candidates in Jurkat cell-based functional assays. Among the five engineered TCRs, all of which demonstrated positive cellular responses, three showed significantly increased activities relative to their templates and one showed substantial enhancement. These functional gains were consistent with more favorable interaction energy from structural and physical modeling. Together, our results support a generalizable paradigm for TCR engineering, in which learned mutation policies can efficiently navigate the peptide-specific binding landscape of TCRs and propose biologically enhanced candidates without explicit structural supervision, offering a practical route for early-stage computational TCR optimization for challenging tumor antigens.

## 1 Introduction

T cells are a critical component of the adaptive immune system. Their activation is triggered by the engagement of cognate T cell receptors (TCRs) with peptide antigens presented by the major histocompatibility complex (MHC). TCRs are heterodimeric proteins whose antigen-specificity is conferred by the *α* and *β* chain; each containing three highly variable and flexible complementary-determining regions (CDRs) 1 − 3 [1]. This modular architecture enables both vast diversity and high context-dependent interaction specificity, which could be leveraged and harnessed to direct the immune system toward pathogenic cells, and in particular, cancer cells. For example, in TCR-T cell therapy, the patient’s own T cells are harvested, genetically modified to express enhanced tumor-reactive TCRs and then infused back into the patient, enabling the immune system to detect and destroy tumor cells expressing the target antigen [2, 3].

Engineering and screening of candidate TCRs with sufficiently strong and specific binding affinity is thus a crucial step and longstanding bottleneck in the process. Because of the close resemblance to self of cancer cells, endogenous tumor-reactive TCRs usually exhibit moderate to low affinity toward tumor antigens (TAs), compared to those recognizing foreign, viral antigens [4–8], as a result of the thymic negative selection to avoid autoimmunity [9]. This reduces their capability in triggering robust tumor rejection, limiting the therapeutic efficacy [10].

To overcome this limitation, several TCR engineering strategies have been developed to enhance the affinity and sensitivity against TAs. High-affinity TCRs could be obtained through affinity maturation [11], directed evolution [12, 13] and gene editing [14, 15], and candidates can be further screened by high-throughput methods including phage display [13, 16–18] and cell-based functional assays [19]. While powerful, large-scale and exhaustive experimental screening still remains expensive and time-consuming.

Computational approaches have therefore been developed to efficiently prioritize candidate TCRs *in silico* from the combinatorial sequence landscape, by predicting the binding specificity between TCRs and peptides. Limited by data availability and labeling noise, most of the existing methods are framed as binary classifications of experimentally-validated true interactions versus non-interacting pairs, and frequently focus on the TCR CDR3*β* (complementarity-determining regions 3 of the *beta* chain) sequence, the most diverse region and a strong empirical determinant for many functional prediction tasks. Several studies have achieved high predictive performance on average across the peptide panel [20–22]. More recent approaches jointly model the other CDR regions such as CDR3*α* to enhance the performance and generalizability [23, 24]. Despite these progresses, computational modeling of TCR-peptide interaction remains challenging due to the intrinsic promiscuity of TCR binding, which coexists with high context-dependent specificity, as well as the substantial structural flexibility of CDR and the limited availability of high-quality training datasets. These factors introduce dataset biases and limit the ability of classifiers to directly guide the rational design of clinically effective TCRs [21, 25].

More recently, generative AI and language models have been applied to the field of protein design [26, 27]. Yet, TCR optimization still poses a distinct challenge, as most of the models focus on the design of isolated proteins without conditioning on their interaction properties and perform poorly on the flexible loop structures that dominate the TCR-pMHC interface. As a result, these models alone are insufficient for capturing the sensitive, multivariate energy landscape of TCR-peptide interactions, while also maintaining biophysical plausibility of the TCR designs. For such tasks, successful design pipelines usually employ an ensemble of multiple specialized modules including backbone design, structural refinement, inverse folding, and functional classifiers, demanding substantial training data and computational resources [28]. This underscores the need for dedicated, domain-calibrated and data-efficient frameworks tailored to the peptide-specific TCR interaction landscape.

In this work, we developed `TCRPPO2`, an integrated, end-to-end framework that leverages a reinforcement learning (RL) algorithm and generative reward models to enable practical, peptide-specific TCR design. Building on our previous work `TCRPPO` [29], which established the advantages of RL over existing approaches for general TCR optimization, `TCRPPO2` advances the paradigm toward biologically actionable design by explicitly resolving the key design bottlenecks: the need for peptide-specific, rational and reliable signals and constraints to ensure the efficacy and viability of the TCR candidates. Specifically, we trained a proximal policy optimization (PPO) RL agent to jointly optimize two complementary objectives: (1) a **peptide-specific binding score** derived from fine-tuned predictive models trained on robustly curated samples integrating biochemically relevant functional annotations; (2) a **sequence validity score** assessed by an unsupervised generative critic trained on large-scale unlabeled TCR repertoires. This dual-objective setup promotes the RL policy to enhance the binding while preserving physicochemical properties within a plausible TCR-like distribution. We evaluated `TCRPPO2` computationally, observing consistent trends of improvement in external evaluations including predictive benchmarking, structural modeling and molecular dynamics simulations. We then validated the approach experimentally using Jurkat cell-based functional assay to measure the reactivity of newly engineered TCRs targeting the clinically relevant MART-1 (Melanoma Antigen Recognized by T cells 1) antigen (ELAGIGILTV), presented by class I MHC HLA-A*02:01. Among five optimized TCRs, all with positive responses in contrast to negative controls, three showed elevated binding affinity compared to the templates, and one with particularly significant enhancement, achieving a 60% rate of successful optimization and 20% rate of significant enhancements. These results demonstrate the competitiveness and feasibility of a purely data-driven, AI-guided computational design-and-screening workflow. Collectively, our work bridges the critical gap between advanced, integrated generative AI models and practical applications in T cell immunotherapy, with potential to significantly accelerate early-stage design and screening of high-affinity candidate TCRs with clinical promise.

## 2 Results

### 2.1 The TCRPPO2 framework for high-affinity TCR design

To systematically address the challenge of task adaptation, data pre-processing and post-selection, we developed `TCRPPO2`, a rigorous, modular, end-to-end framework that could be flexibly deployed for peptide-specific TCR optimization and screening (Fig. 1B). Central to `TCRPPO2` is an RL method that learns a step-wise mutation policy to enhance the binding affinity of TCRs toward target peptides [29]. The **agent** predicts a policy that takes actions (mutations introduced to the input sequence) given the current sequence, which are then independently evaluated by the **environment** with two main objectives (4.1.3):

- **Peptide-specific binding score**: the predicted binding affinity score between the obtained sequence and the target peptide from a pre-trained, frozen classifier model based on the Attentive Variational Information Bottleneck (AVIB) architecture [30] (4.2.1).
- **Sequence validity score**: whether the sequence falls within the distribution of naturally-occurring TCRs, indicating its plausibility and synthesizability, evaluated by an independent generative critic trained on large-scale unlabeled TCR sequence corpus (4.2.2).

**Fig. 1.**
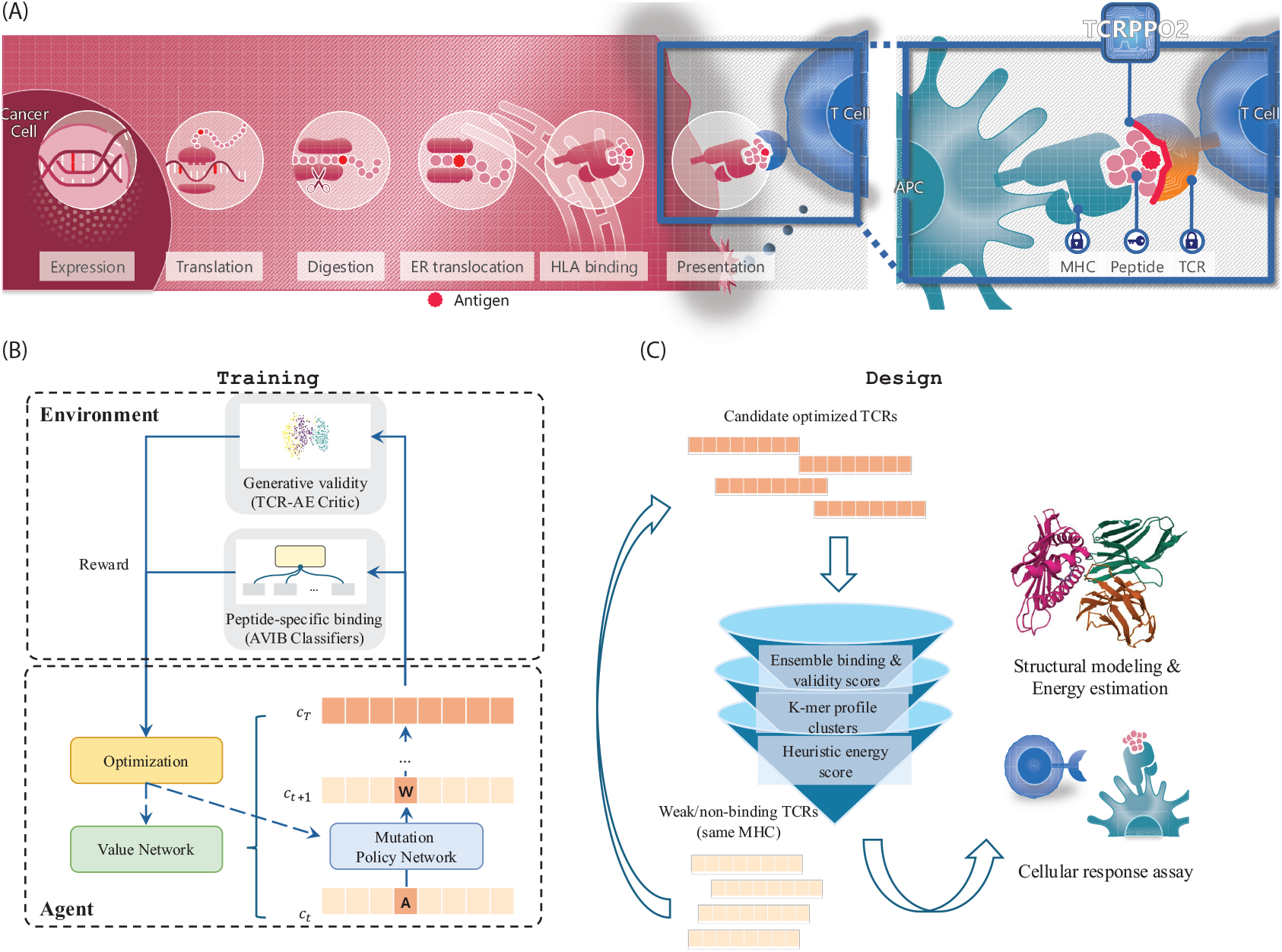
The TCRPPO2 framework. (A) The antigen presentation and T cell recognition pathway. Cancer-specific antigens are expressed and presented by MHC (HLA) proteins on the cell surface, then recognized by specific TCRs on T cells to trigger downstream immune response. The `TCRPPO2` framework computationally optimizes the interaction between TCRs and peptide-MHC complexes; (B) Schematic of the reinforcement learning (RL) framework of `TCRPPO2` : the agent proposes and executes mutations given the input TCR and peptide sequences, which are evaluated by the environment composed of a generative critic assessing sequence validity and an ensemble classifier estimating peptide-specific binding affinity; (C) Overview of the complete data processing and optimization workflow of `TCRPPO2` : weak or non-binding TCRs are input to the RL framework to obtain optimized candidates, which then undergo successive fast computational screening, followed by validation through structural modeling and cell-based functional assays.

The objective function rewards mutations that lead to both higher binding affinity and validity, and their feedback is then back-propagated to update the policy network. This setup encourages the agent to explore the mutation space and learn an optimal mutation policy that cumulatively mutates the given sequence to achieve the highest rewards, balancing higher binding affinity and adherence to the distribution of naturally occurring TCRs (see 4.1 for more details). The policy model could then be directly applied to template TCR sequences to optimize their binding affinity toward the given peptide.

We tailored the RL training and post-generation filtering framework to achieve more robust peptide-specific designs in each stage of the `TCRPPO2` pipeline (Fig. 1):

- **Specification of the AVIB classifier and the** `TCRPPO` **model:** Our pipeline finetunes both the AVIB binding affinity classifier and the `TCRPPO` RL model specifically for the given target peptide, to overcome the huge imbalance across peptides and the heterogeneity of the positive/negative space in generic training data derived from VDJdb [31], McPAS [32] and IEDB [33].
- **MHC-restricted data selection and split:** The selection of the negative training samples for the classifier and the RL model, as well as the template sequences for the optimization experiments, were restricted to TCRs that recognized alternative peptides presented by the same MHC. This strategy is employed to prevent the models from relying on overly simplistic or trivial patterns, and avoid optimization tasks that are infeasible or irrelevant.
- **Post-selection:** To provide external computational screening for biochemically viable designs, we employ a layered filtering approach including several orthogonal heuristics to efficiently narrow down the thousands of generated TCR sequences from `TCRPPO` outputs. We first use k-mer-based sequence clustering and Miyazawa-Jernigan binding energy estimates [34] for fast screening. For more fine-grained selection, we perform more accurate yet computationally demanding structural prediction, molecular dynamics and energy estimation to further validate the candidates.

In the following sections, we present a detailed description of the `TCRPPO2` framework, computational benchmarking on more than 2,000 TCR templates, and the results of practical optimizations of selected weak binders accompanied by experimental validation.

### 2.2 Benchmarking of TCRPPO2 on peptide-specific datasets

Most existing TCR–pMHC prediction methods emphasize aggregate performance across diverse peptide repertoires, which may bias models toward more prevalent peptides while limiting their ability to accurately predict interactions involving rarer or highly specific epitopes. To enhance the applicability toward the specific peptides of interest, we tuned the AVIB classifier [30] using either AVIB’s own embedder or embeddings from fine-tuned ProtBERT [35], on peptide-specific data. Specifically, we used the *beta*-only TChard dataset [25] and only selected samples related to the target peptide (MART-1 epitope ELAGIGILTV). Each model was trained and tested on the same five random data splits. On average, the fine-tuned models achieve higher performance by AUROC than the generic model, and the ensemble of the fine-tuned models achieve the highest performance as well as less variance across data splits (Fig. 2A).

**Fig. 2.**
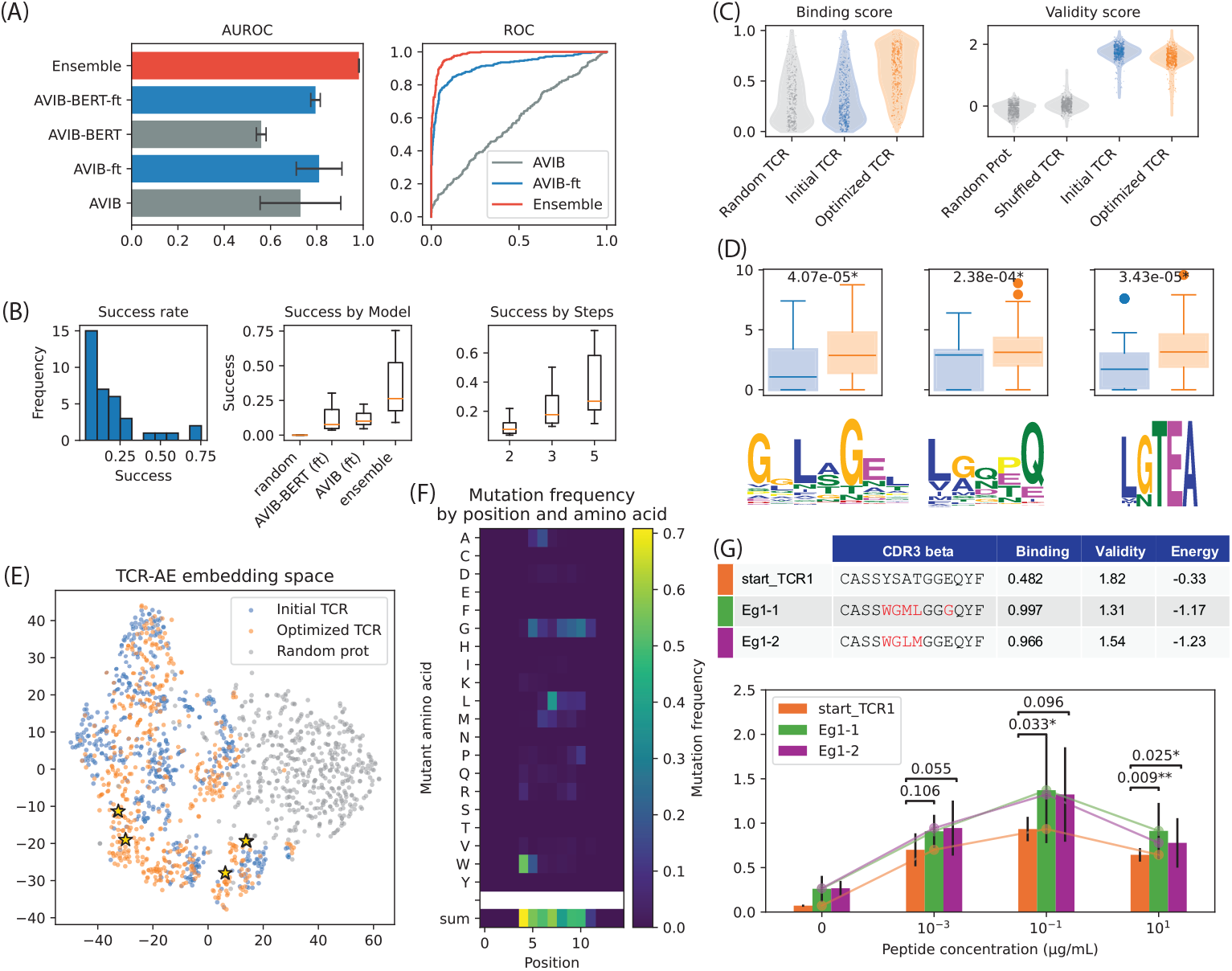
Experimental results on MART-1-binding TCR optimization. (A) Performance of baseline, fine-tuned and ensemble AVIB classifiers on MART-1 binding prediction, across five random splits; (B) Left: success rate (binding score *>* 0.9, validity score *>* 1.2577) distribution; Middle: success rate by the binding score reward model; Right: success rate by the number of mutation steps; (C) Benchmarking of the binding score (left) and validity score (right) between initial and optimized TCR sequences and random controls; (D) The enrichment of binding-related sequence motifs in the optimized TCRs compared to the initial TCRs; (E) The distribution of initial and optimized TCRs and random controls in the latent embedding space of the TCR-AE; the stars denote the experimentally validated TCR designs in Fig. 2G and 3F; (F) The mutation frequency by position, aggregated across all optimized TCRs of 15 residues length; (G) The reporter cell assay results for the selected initial (start TCR1) and optimized TCRs (Eg1-1 and Eg1-2).

We then trained `TCRPPO2` on a selected set of starting TCRs that recognize peptides presented by the same MHC variant HLA-A*02:01 from VDJdb [31] with 3 or 5 maximum modification steps. This is to exclude TCR templates whose binding modes differ substantially and would compromise the feasibility of the optimization. The trained policy models were applied to 2,831 selected TCR templates from VDJdb by the same selection criteria with 2, 3 and 5 maximum modification steps. Using a random mutation policy as the baseline, we demonstrate that the learned `TCRPPO` policies achieve a substantially higher success rate (binding score *>* 0.9, validity *>* 1.2577; Fig. 2B). As expected, using the ensemble classifiers as the binding affinity scoring function achieves the highest performance. Furthermore, we observe that increasing the number of mutation steps further improves the success rate, with 5 mutation steps achieving success rates ranging from 0.12 to 0.75 (average 0.37) across models. These results indicate that the search for optimal mutations is a non-trivial task, necessitating the use of the policy model to effectively navigate the vast mutation space (∼ 600 million possibilities for one template sequence of length 10).

Specifically, we show that the optimized TCRs achieve significantly higher predicted binding score than randomly selected TCRs and their initial TCR templates (Fig. 2C left), while preserving a similar validity score distinguishable from invalid control sequences such as random protein segments and shuffled TCR sequences (Fig. 2C right). This highlights the effectiveness of the learned RL policies in jointly optimizing both objectives. Moreover, several sequential motifs enriched in the positive training set also attain higher scores in the optimized TCRs relative to the templates (Fig. 2D). Both optimized and initial TCRs were indistinguishable in the embedding space of the TCR-AE critic model, further demonstrating the effectiveness of the validity objective (Fig. 2E). By investigating the behavior of the mutation policies, we observe a tendency of the policy model to introduce hydrophobic and non-polar residues (leucine, methionine, tryptophan, glycine; Fig. 2F), which aligns with previous discoveries that TCR-pMHC interactions favor hydrophobic interactions within the interface [36–38]. Our analysis implies that the `TCRPPO2` policy model successfully transfers biologically meaningful binding-related sequential patterns from the training data to the optimization outputs, and achieves robust computational performance on a broad range of TCR templates.

### 2.3 *In silico* design of MART-1 epitope-binding TCRs

In real-world applications, it is usually more practical to optimize existing low-affinity TCRs with known weak or medium binding to preserve the stability and the valid binding pose and surface [10], as in affinity maturation experiments [7, 39]. Following this established principle, we selected template TCRs from existing databases with full sequence information and weak binding confirmed by both computational prediction and experimental evidence. We ran trained `TCRPPO2` models on the selected templates, pooled the results then ranked and screened with the same heuristics as in 2.2. For experimental validation of the binding, the optimized TCRs were expressed in Jurkat cells (human T cell leukemia line) and co-cultured with antigen-presenting cells (APCs) with increasing peptide concentration from 0 to 10^1^*µ*g*/*mL (details in 4.6).

#### 2.3.1 Optimized weak MART-1 binders show experimentally validated affinity enhancement

We ran `TCRPPO2` on a candidate template TCR (CASSYSATGGEQYF) derived from the TCR repertoire of a melanoma patient [40] with ensemble classifier score = 0.48, indicating borderline binding. We then selected two engineering results with *>* 0.95 binding scores and top-ranked Miyazawa-Jernigan energy (Fig. 2G). Both candidates are produced by the ensemble AVIB predictor as the reward function.

The optimized TCRs along with the template were then tested in Jurkat reporter cell assays. Successful TCR expression in the Jurkat cells was confirmed by flow cytometry (Fig. S3 ∼ S5). We observed that both optimized TCRs exhibit enhanced antigen-specific T cell activity compared to the template at higher dosages, leading to a near 100% hit rate (Fig. 2G). The outcomes align closely with the binding prediction and the energy-based heuristics, indicating that the `TCRPPO2` policy model effectively optimizes the template TCR for peptide recognition.

#### 2.3.2 Knowledge-guided designs strengthen TCR optimization results

Having preliminarily established the biological relevance of our learned `TCRPPO2` policy model, we next aimed to further improve its accuracy and practical utility. To this end, we refined both the curation of the training data and the selection of templates, incorporating prior biological knowledge to more effectively capture the landscape of binding affinities. In particular, we observed that the predictive model trained directly on binary TCR-peptide interaction pairs fails to adequately capture the “intermediate” interactions identified by IEDB [33], which have the assay label “multimer/tetramer qualitative binding; Positive-Intermediate” (Fig. 4). Such cases, labeled as positive in the training data, would lead to misclassifications of similar intermediate or weak binders as positive during evaluation. To address this issue, we excluded the intermediate pairs and re-trained the model using only verified strong binders and non-binders. The resulting model showed both higher performance (Fig. 3A) than the baseline models and also more consistent ranking of the “intermediate” interactions in IEDB, correctly placing them between the experimentally validated true positives and negatives (Fig. 3B). This indicates that the model is capable of interpolating between the sequential patterns of strong binders and non-binders to weak binders, thereby offering more biologically meaningful guidance to the RL-based modeling.

**Fig. 3.**
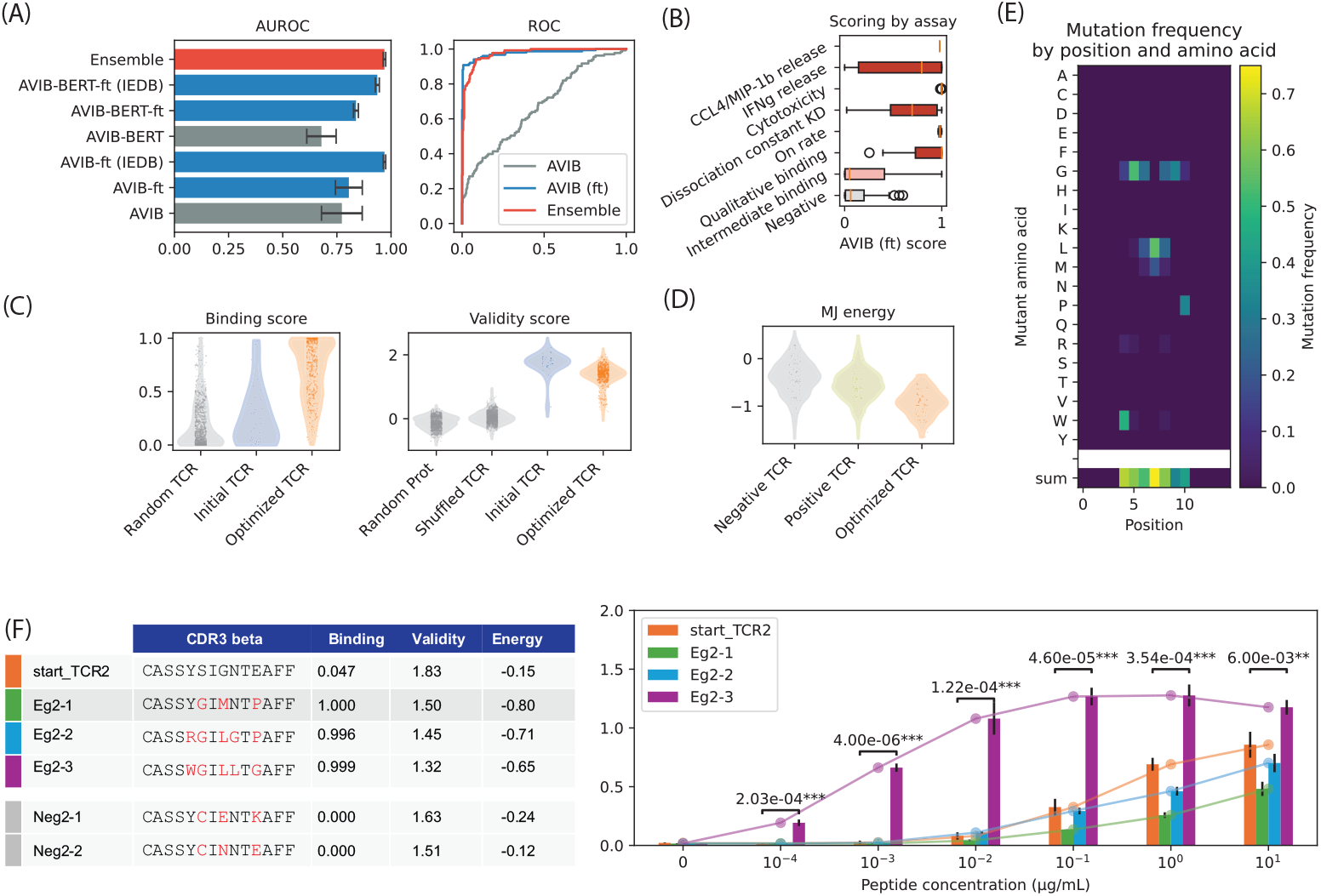
Experimental results on MART-1-binding TCR optimization with knowledge-guided designs. (A) Performance of baseline, fine-tuned and ensemble AVIB classifiers with or without IEDB annotation-guided sanitization on MART-1 binding prediction, across five random splits; (B) Predictions of the sanitized AVIB classifier on IEDB test samples, stratified by experimentally validated annotation labels; (C) The comparison of the binding score (left) and validity score (right) between initial and optimized TCR sequences and random controls; (D) The comparison of the Miyazawa-Jernigan interaction energy with the MART-1 epitope between initial and optimized TCR sequences and random controls; (E) The mutation frequency by position, aggregated across all optimized TCRs of 15 residues length; (F) The reporter cell assay results for the selected initial (start TCR2) and optimized TCRs (Eg2-1, Eg2-2 and Eg2-3); the negative controls have near-zero response on all concentration and are thus omitted.

**Fig. 4.**
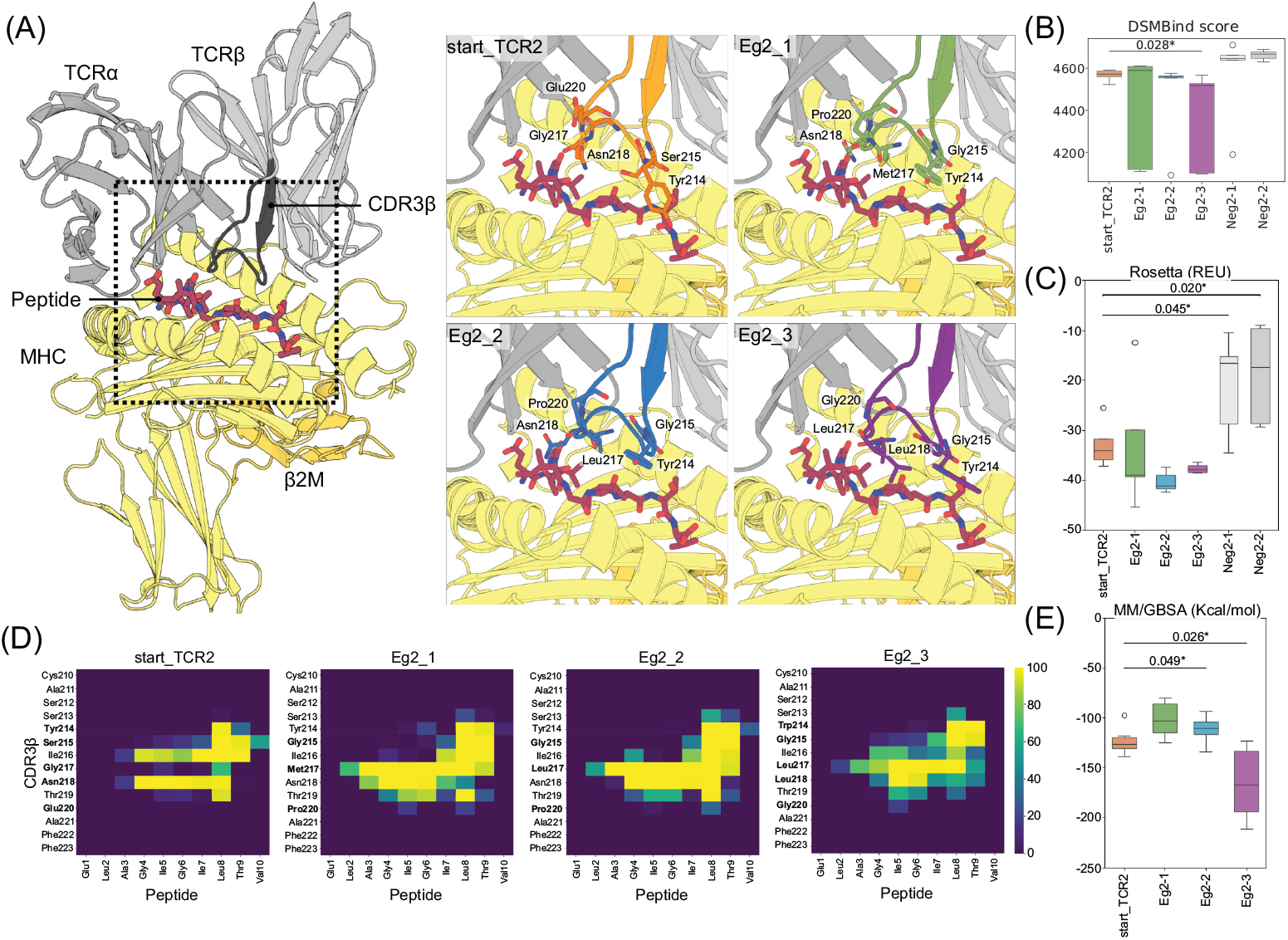
Physical modeling and energy estimation results. (A) The predicted TCR-pMHC complex structures for start TCR2, Eg2-1, Eg2-2 and Eg2-3; (B) DSMBind energy estimates (Mann-Whitney U test); (C) Rosetta energy estimates; (D) Interface contact matrices from MM/GBSA; (E) Binding affinity estimates from MM/GBSA.

We used this “sanitized” AVIB classifier as the reward to train `TCRPPO2`, then ran the optimization pipeline with the held-out intermediate binders from IEDB as templates. We observed sharper increases in the predicted binding score of the optimized TCRs compared to the templates and an adequate preservation of validity (Fig. 3C). Our optimized TCRs also display significantly lower Miyazawa-Jernigan energy, which favored average positive samples over negatives (Fig. 3D), as well as more focused mutation preferences (Fig. 3E).

We selected one of the “intermediate binding” templates with low predicted binding score and experimentally characterized three top-ranked optimized TCRs with the lowest Miyazawa-Jernigan energy and diverse mutation profiles, as well as two negative controls with random mutations on the same sites and low binding scores. Among the three positive TCRs, one (Eg2-3) exhibited significantly increased binding affinity at all concentrations compared to the template, while the other two maintained a lower but comparable response. In contrast, neither of the negatives displayed measurable binding (Fig. 3F, Fig. S2).

These results demonstrate that our method could be applied to the more nuanced task of fine-tuning TCR binding affinity, and that the heuristic computational screening of the starting templates can further enhance the performance. Combined with the previous round of experiments, we attained a 3*/*5 hit rate in a mutational space of around 1 × 10^8^ possibilities, while successfully discriminating against deleterious mutations, further supporting `TCRPPO2`’s capability in exploring the interaction landscape of TCR sequences and yielding optimal mutation policies that are difficult to achieve through conventional strategies.

### 2.4 Structure and energy prediction of optimized TCRs

`TCRPPO2` generates a diverse ensemble of candidate TCR sequences for a given peptide-MHC (pMHC) target, necessitating quantitative criteria to prioritize designs for experimental validation. To this end, we first employed TCRmodel2 to generate structural models for the reference TCR (start TCR2), three engineered variants (Eg2-1, Eg2-2, Eg2-3) and the two negative controls (Neg2-1, Neg2-2) bound to the pMHC complexes (Fig. 4A). Next, we applied DSMBind [41], an unsupervised binding energy prediction framework, fine-tuned on the TCR-pMHC structural data, to score the predicted structures of our samples (Fig. 4B). The selected positive TCRs generally exhibit significantly lower and more negative binding energy than the negative controls (Mann-Whitney U test), suggesting a consistency between our data-driven objective and external energy estimates.

Next, we implemented a physics-based computational pipeline encompassing Rosetta software suite for estimating binding energies [42] and excluding inactive TCR candidates, followed by molecular dynamics (MD) simulations and molecular mechanics-generalized Born surface area (MM/GBSA) calculations to rank the putative active designs (Fig.S12). The five top-ranked TCR models for each complex generated by `TCRPPO2` were ranked using the Rosetta energy scoring with backrub sampling, and the average energy across models was used for ranking. (Tab. S2 metrics from TCRmodel2). The Rosetta scoring function identified the negative controls as the highest-energy complexes, in line with the results obtained with DSMBind (Fig. 4C).

To capture dynamic fluctuations, we performed conventional MD simulations combined with MM/GBSA analysis. For each system, the two top-scoring TCRmodel2 [43] models were refined by adding the missing *β*-microglobulin (*β*2m), MHC *β*-chain, and associated N-glycans [44], producing biologically realistic TCR-pMHC complexes (Fig. 4). Each refined complex was subjected to 120 ns of MD simulations per system across six independent replicas. To ensure that the simulations reached equilibrium and provided a reliable basis for the MM/GBSA calculations, we analyzed the structural stability and flexibility of each complex. The root-mean-square deviation (RMSD) of C*α* atoms indicated structural convergence within 2 °A for all systems (Fig. S13). Residue-level root-mean-square fluctuation (RMSF) profiles revealed no major differences across complexes, although the MHC *β*-chain in Eg2-1 displayed increased flexibility, reflecting local conformational adjustments without affecting overall complex integrity (Fig. S14). These results indicate that the introduced sequence modifications do not substantially perturb the intrinsic dynamics of the TCR-pMHC interface. To complement the global stability analysis, we examined residue-level contacts between TCR and pMHC to identify specific interactions underpinning complex stability. Contact matrices revealed alterations extending beyond the TCR-pMHC interface, indicating that CDR3*β* mutations can induce long-range structural rearrangements (Fig. S15). Enlargement on the interface between CDR3*β* and peptide highlighted broader contacts in the Eg2-3 interface, consistent with a more compact and stable binding interface (Fig. 4C). These analyses provide a mechanistic view of mutation-dependent changes in interface organization and their contribution to complex stability.

Finally, binding affinities of the TCR-pMHC complexes were estimated using MM/GBSA calculations following the protocol of Crean et al [45]. MM/GBSA indicated a significant decrease in binding affinity for Eg2-1 relative to the reference system, whereas Eg2-2 showed no significant change, consistent with comparable activity. Eg2-3, corresponding to the TCR with the highest experimental activity, exhibited the strongest predicted affinity, in agreement with experimental observations (Fig. 4D, E), providing a reliable and consistent ranking of the engineered TCRs.

## 3 Discussions

In this study, we present and experimentally validate `TCRPPO2`, an end-to-end, automated reinforcement learning framework for TCR optimization that integrates robust pre-processing, data-driven generation and knowledge-guided post-selection. The framework forms a coherent pipeline that directly addresses several persistent limitations in generic protein sequence design workflows, including imbalanced training data, unrealistic template selection, invalid or low-quality outputs, and insufficient incorporation of biochemical constraints that often hampers practicality. By combining data-driven deep reinforcement learning (RL) models with a learned generative critic that evaluates and shapes the candidate sequences, `TCRPPO2` efficiently optimizes low- or moderate-affinity TCR templates into high-affinity variants. Importantly, this is achieved with minimal reliance on computationally expensive physics-based simulations or labor-intensive experimental screening. The resulting TCR sequences are not only diverse but also biologically plausible, effectively capturing the learnable “positive” sequence patterns from the training data, leading to meaningful improvements in peptide-specific affinity, further supported by external validations such as motif enrichment analyses and interaction energy heuristics. The generative critic also serves as a regulatory mechanism that prevents major deviations from the natural TCR sequence distributions, thereby improving practical likelihood of synthesis and expression. We further strengthened the pipeline through knowledge-based post-selection, such as sequence clustering and rapid interaction energy filters, to prioritize candidates that are not only computationally promising but also mechanistically plausible. This layered design substantially enhances reliability at each stage by grounding exploratory generation in relevant biochemical constraints. Reporter cell assay results confirmed that our computationally optimized TCRs exhibited strong cellular activity without requiring preliminary experimental screening, consistent with post hoc physical modeling results that indicated improved binding energies. Collectively, these findings indicate that `TCRPPO2` offers a flexible, rigorous and data-efficient framework for practical TCR engineering. By tightly integrating RL-driven exploration with bio-chemically informed priors, `TCRPPO2` efficiently leverages limited interaction data to navigate the vast TCR mutation space and provides a scalable strategy for identifying high-affinity, high-quality candidate TCRs with potential translational relevance.

We implemented a physics-based modeling pipeline aimed at ranking the ensemble of candidate TCR sequences for TCR-pMHC complexes as well as to investigate the structural and energetic consequences of the TCR mutations. Initial binding estimates were obtained using DSMBind, an unsupervised TCR-pMHC scoring model, and Rosetta with backrub sampling, which explores local backbone flexibility. Variants lacking measurable experimental activity tended to show unfavorable predicted binding energies, whereas active engineered TCRs exhibited a broader range of scores. These results indicate that, while DSMBind and Rosetta predictions do not quantitatively capture functional activity, they provide an initial filter to exclude likely inactive sequences, complementing experimental characterization. To gain atomistic insight, we performed MD simulations on complete TCR-pMHC complexes, enabling evaluation of structural stability and residue-level interactions. RMSD and RMSF analyses confirmed that global complex architectures were largely preserved across variants, while contact analyses revealed that CDR3*β* mutations could trigger distal structural rearrangements, as previously suggested [46], modulating interactions beyond the binding interface. Notably, the Eg2-3 variant, which exhibited the strongest response in the cell-based functional assays, showed broader CDR3*β*–peptide contacts and a more compact binding interface, suggesting that targeted mutations can enhance specificity and stability without compromising overall complex integrity. Binding affinities estimated via MM/GBSA reflected these structural trends and provided a physically grounded, ensemble-averaged measure of relative ΔΔG. MM/GBSA combines conformational sampling from MD with implicit solvation energy calculations, capturing dynamic fluctuations and residue-level contributions that static scoring functions cannot [45]. The resulting ranking was broadly consistent with experimental activity measurements and offered insight into which interactions contribute most to binding. These combined computational approaches provide complementary perspectives on TCR-pMHC recognition, though it is important to note that binding affinity alone does not fully determine T cell functionality, which also depends on kinetic parameters, co-receptor engagement, and cellular context [47, 48].

Most of the existing TCR-peptide interaction datasets, including those used to train `TCRPPO2`, reduce “peptide recognition” to a binary outcome label that aggregates results from heterogeneous experimental modalities against a restricted set of test peptides, such as binding assays (multimer/tetramer staining), and *in vitro* functional readouts [31]. Because these modalities capture distinct aspects of TCR-peptide interactions, models trained directly on such aggregated labels risk conflating multiple underlying biological determinants. In addition, many datasets under-represent interaction kinetics (association/dissociation rates), which govern both the rate and the stability of the complex formation, and have well-established functional consequences [49]. For example, in the case of the MART-1 epitope analyzed in this study, most available measurements derive from multimer-based assays, potentially biasing the model toward features associated with avidity rather than monomeric binding affinity, which are partially related yet not equivalent. Moreover, extremely high-affinity or high-avidity TCRs can undergo negative regulation, as a safeguard against autoimmunity [50, 51]. Several studies have suggested an optimal affinity window with an “upper bound” required for reliable antitumor immune responses [52]. Despite these limitations, we have shown that with appropriate data curation guided by biological priors, the output probabilities of the AVIB classifier trained on binary labels could approximate the distinctions between weak and strong binders, effectively providing a coarse surrogate estimation for more granular binding affinity measurements. Nevertheless, addressing this issue more rigorously will require moving beyond exclusive reliance on binary labels. Our framework could be further strengthened by leveraging continuous energy-based predictions as reward functions, thereby enabling finer-grained control of TCR-peptide binding affinity. The results from computational structural prediction [43] and unsupervised energy scoring from DSMBind [41] underscore the promise of this direction for more precise modeling of TCR-peptide interactions.

In real-world immunotherapeutic applications, computational TCR design frameworks must holistically evaluate and jointly optimize all the aforementioned intrinsic biochemical and biophysical properties (e.g. affinity, avidity, kinetics) together with potential adverse effects such as off-target or on-target-off-tumor reactivity. Therefore, to achieve clinically relevant TCR designs, these factors should be independently modeled and mechanistically disentangled to support precise controls along each functional axis. Given current limitations in data availability and the substantial computational cost of detailed physical simulations, progress will depend on both continued accumulation of high-quality experimental data and the development of more efficient predictive methods. This challenge also underscores another particularly compelling advantage of reinforcement learning: its capacity to integrate multiple, potentially non-differentiable reward functions, allowing the policy model to navigate the complex multi-objective landscapes and achieve biologically meaningful trade-offs, such as improving binding affinity while balancing cross-reactivity and off-target effects, ultimately facilitating the design of TCRs with improved efficacy and enhanced safety.

Recent advances in large protein language models (PLMs) [53, 54] further establish reinforcement learning as a powerful approach for fine-grained control in sequence generation and functional optimization. By harnessing the generative capacity of the PLMs and aligning them with biologically grounded objectives, our proposed framework could be scaled to incorporate richer evolutionary and structural priors, facilitating the rational design of TCRs with enhanced affinity alongside high sequential and structural validity. Concurrent progress in computational structure predictions of proteins and protein-protein interaction complexes also opens opportunities for direct optimization in the 3D structure space even for previously uncharacterized interactions. Such capabilities support more stringent sequence-structure co-design with greater scale, precision, and awareness of the underlying physicochemical and geometrical constraints.

In summary, our work demonstrates the effectiveness of data-driven reinforcement learning for the rational optimization of structurally and functionally relevant TCRs. The flexibility of this approach, including its ability to accommodate multiple design objectives and larger-scale modeling, positions it as a powerful and versatile platform for future explorations in TCR engineering and more broadly functional protein design.

## 4 Materials and Methods

### 4.1 TCRPPO2

#### 4.1.1 Problem definition

Let *c* be a CDR3*β* sequence of a TCR and *p* be the peptide. In the following sections, we use “CDR3*β*” and “TCR” interchangeably unless specified. Based on the framework of `TCRPPO` [29], we formulated the problem of TCR optimization as a step-wise Markov Decision Process (MDP) that (1) improves binding toward the peptide *r*_*b*_ and (2) maintains or improves sequence validity *r*_*v*_ of a template TCR sequence: ℳ = {𝒮, 𝒜, 𝒫, ℛ} with the following components:

- The **state** space 𝒮: the state at each step *t* is defined as a tuple *s*_*t*_ = (*c*_*t*_, *p*) containing the current TCR sequence and the peptide sequence. We denote *s*_0_ as the input state and *s*_*T*_ as the final state that either satisfies both objectives or hits the maximum exploration steps.
- The **action** space 𝒜: an action determines the changes introduced to the state at each step. In our setting, the action **a** is a choice of mutation that determines both the mutation site *i* and the mutant residue type *o*.
- The state transition probability 𝒫: in our setting, the transition probability into a new state (updated TCR sequence) given the current state (input TCR sequence) and action (mutation) is deterministic, i.e., 𝒫(*s*_*t*+1_|*s*_*t*_, **a**_*t*_) = 1
- The **reward** ℛ: the evaluation of every state, obtained from the generative critic of *r*_*v*_ and the predictive model of *r*_*b*_ (see 4.1.3 for more details)

Based on this formulation, the optimization process becomes sequential decision-making: given a starting state *c*_0_ = (*s*_0_, *p*) with a template TCR sequence *c*_0_ and a target peptide *p*, an RL agent navigates the combinatorial protein sequence space by iteratively introducing mutations to the current sequence, which resembles the natural process of protein evolution or affinity maturation. The objective of the training is to learn an optimized policy *π*(**a**|*s*), which determines the mutation action **a**_*t*_ from the state *s*_*t*_, to maximize the expected reward of the output sequence *s*_*T*_ at the final step *T* .

#### 4.1.2 Mutation policy

At any given time step *t*, the policy *π*(**a**|*s*) predicts an action **a**_*t*_ = (*i*_*t*_, *o*_*t*_) that specifies a position *i*_*t*_ to introduce the mutation and a residue type *o*_*t*_ as the mutant. Specifically, each residue *c*_*i*_ in sequence *c* is first embedded as a concatenation of the BLOSUM vector, one-hot encoding and a learnable embedding, then provided to a bi-directional LSTM [55] to obtain the latent embedding:

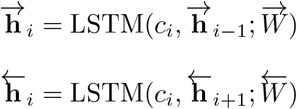

where 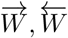 are the parameters of the two LSTM directions. The embedding of each position is concatenated from the respective embeddings of both directions:

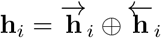

The full sequence-wise embedding is concatenated from the endpoint embedding of each direction:

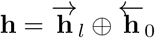

where *l* is the length of the sequence.

Let **h**_*t*_ and **h**_*p*_ be the full sequence embedding of the TCR sequence *c*_*t*_ and the peptide sequence *p*, respectively, and **h**_*i,t*_ is the embedding of the *i*-th position of *c*_*t*_. The probability of choosing position *i* for the mutation is defined as a softmax over all positions:

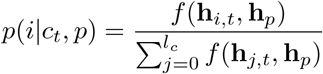

where the function *f* is a learnable multi-layer perceptron with two layers and rectified linear unit (ReLU) activation.

Given *i*, the choice of mutant *o* is then defined as:

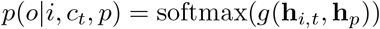

where the function *g* is a learnable multi-layer perceptron with two layers and rectified linear unit (ReLU) activation that projects the embeddings **h**_*i,t*_ and **h**_*p*_ into a 20-dimensional vector each corresponding to a residue type. To balance exploration and exploitation, both *i*_*t*_ and *o*_*t*_ are sampled from the predicted probability distribution.

#### 4.1.3 Reward function

The reward function is defined with two components: a peptide-specific binding score and a sequence validity score. Specifically, given any exploration trajectory (*s*_0_, **a**_0_, *s*_1_, **a**_1_, …, *s*_*T* −1_, **a**_*T* −1_, *s*_*T*_), where each state *s*_*t*_ is the result of applying action **a**_*t−*1_ on the previous state *s*_*t−*1_, the outcome is evaluated by the score of the final state *s*_*T*_ with the following scoring functions:

##### Binding score

The binding score *r*_*b*_(*c*_*T*_, *p*) is predicted from the TCR and peptide sequences using any trained AVIB classifier from 4.2.1, or their ensemble.

##### Validity score

*r*_*v*_ evaluates whether the sequence *c*_*t*_ has similar sequential patterns as naturally occurring TCRs, which we formulate as an anomaly detection problem using generative models [56, 57], where a generative model estimates how well the sample aligns with the latent space derived from real samples. Following the practices in [29, 58], we use an autoencoder trained on an independent set of 277 million unlabeled sequences from TCRdb [59], denoted as `TCR-AE`. The validity score of any sequence *c* is then defined by the reconstruction accuracy and the likelihood within the `TCR-AE` latent space:

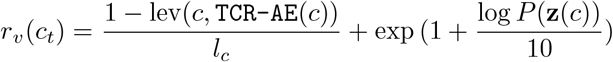

where `TCR-AE`(*c*) is the reconstruction of *c* by `TCR-AE`, *l*_*c*_ is the length of *c*, lev refers to the Levenshtein distance, **z**(*c*) is the latent embedding of *c* by `TCR-AE`(*c*), and the log-likelihood function *P* is defined by a Gaussian mixture model (GMM) of `TCR-AE`’s latent space with *K* = 8 components:

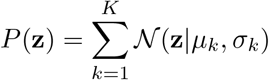

The final **reward function** is calculated from the two scores as an estimate of the projected outcome of each step *t* in the trajectory:

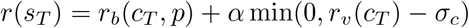

where *α* = 0.5 is a regularization hyperparameter and *σ*_*c*_ = 1.2577 is the threshold for validity following [29].

#### 4.1.4 Proximal policy optimization

Assume the policy *π* is parametrized by Θ. We used the proximal policy optimization (PPO) [60] to learn the optimal policy *π* that maximizes the future rewards:

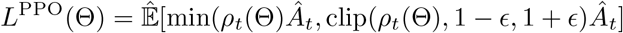

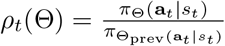 is the ratio between the probability of choosing action **a**_*t*_ with the current and the previous policy, clipped by interval [1 − *ϵ*, 1 + *ϵ*].

*Â*_*t*_ is the estimated advantage of the selected action **a**_*t*_:

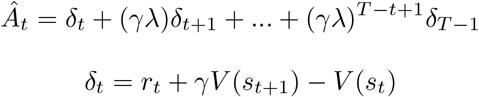

where *γ* ∈ (0, 1) is the discount factor and *λ* ∈ (0, 1) is a regularization hyperparameter, *r*_*t*_ is the reward on the intermediate state *s*_*t*_ (*r*_*t*_ = 0 if *t* ≠ *T* in our setting). *V* (*s*_*t*_) is the value model that predicts the discounted reward-to-go of the current state *s*_*t*_. Let Θ_*V*_ be the parameters of *V*, the training objective for *V* is to :

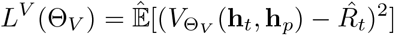

where 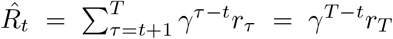 is the discounted reward-to-go. The final objective function is to minimize:

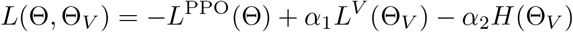

where *α*_1_ and *α*_2_ are weighting hyperparameters and the last term is an entropy regularization for the model parameters [60].

### 4.2 Model implementation and training

#### 4.2.1 Training of the TCR-peptide binding score model

The **AVIB** [30] model receives as input a sample consisted of sequence representation *X*, in our case residue-wise BLOSUM50 encoding [61] of the TCR and the peptide, to learn a compressed latent representation *Z* that predicts the target label (binding) *Y* through the variational information bottleneck (VIB) mechanism:

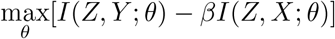

where *θ* is the model parameter and *β* is the regularization hyperparameter. This objective ensures that *Z* preserves only maximal information about the binding *Y* and minimal irrelevant information. Experimental results show AVIB is able to reliably classify out-of-distribution samples, supporting its use as a reward function that stabilizes the RL agent’s exploration process.

We collected TCR-peptide interaction data compiled by Grazioli et al [25] and selected MART-1-specific samples for the training of the AVIB binding classifier prediction. For the data sanitization, samples that overlapped with IEDB [33] entries with “multimer/tetramer qualitative binding; positive-intermediate” assay outcomes were considered weak binders and excluded from the training data. We used the AVIB model trained on general TCR-peptide interaction data, and fine-tuned it on the MART-1-specific subset with learning rate 1 × 10^−4^, batch size 128 and regularization parameter *β* = 1 × 10^−6^.

To experiment with more expressive sequence representations, we implemented the **AVIB-BERT** variant. We first fine-tuned a ProtBERT [35] model on CDR sequence data with learning rate 1 × 10^−6^ and batch size 32. We then provided the fine-tuned embedding of the TCR and the pre-trained embedding of the peptide as inputs to the AVIB classifier and trained the model with the same setup.

#### 4.2.2 Training of the generative critic model

The critic model is an autoencoder with 16 hidden layers and a latent space dimension of 64. The autoencoder is trained on the ∼ 277 million unlabeled TCR CDR3*β* dataset from TCRdb [59] with batch size 256 for 100, 000 steps.

#### 4.2.3 Training of the reinforcement learning framework

To construct a meaningful sequence space for the training of `TCRPPO2`, we provided all TCR sequences from VDJdb [31] with complete annotations (V and J gene usage for both *α* and *β* chains), resulting in 5, 088 unique sequences, then split into training and test data by 4 : 1. During each training iteration, a sequence is first sampled from the training set, mutated incrementally by the current policy, then the final state *s*_*T*_ is evaluated with the reward function. The full trajectory (*s*_0_, **a**_0_, *s*_1_, **a**_1_, …, *s*_*T* −1_, **a**_*T* −1_, *s*_*T*_) and reward *r*_*T*_ are then used as one sample to update the policy and value models through back-propagation.

A complete list of hyperparameters can be found in Supplementary Table S1. Training and inference were conducted on four RTX3090 GPUs. We trained multiple model variants by alternating the configurations of the binding score estimator, maximal mutation steps and training step as in Table 1. Specifically, each variant was trained for 1,000,000 steps and checkpoints at 200,000, 500,000 and 1,000,000 steps were retained for the downstream experiments.

**Table 1.** Variants of TCRPPO2 configurations used for downstream experiments.

| Hyperparameter | Value(s) |
| --- | --- |
| Maximum mutations | 2, 3, 5 |
| Fine-tuning set | full, IEDB-sanitized |
| Fine-tuning split | 1 – 5 |
| Binding score model | AVIB, AVIB-BERT, Ensemble |

### 4.3 Computational TCR optimization

The test samples for the optimization experiments were high-confidence weak binders carefully selected from two sources: (1)The TCR3d [62] samples with high experimentally measured *K*_*D*_ (*>* 10*µM*); (2) The binders from “multimer/tetramer qualitative binding; positive-intermediate” assays in IEDB. Both sources were further screened and ranked by predicted binding scores from the ensemble AVIB predictors from 4.2.1 and those with lowest scores were used for experimental validation. For each template TCR, we ran all the aforementioned policy model variants from 4.2.3 with 2, 3 and 5 maximal mutation steps.

#### 4.3.1 Post-filtering

We pooled and de-duplicated the results from all TCR templates, model variants and mutation steps, and then applied the following screening procedure:

- **Predicted scoring:** We retained sequences with generative critic validity score *>* 1.25 and ensemble AVIB classifier binding score *>* 0.8. For the count of “success” cases in Section 2.2, we specifically only evaluated on templates with initial binding score *<* 0.2. For the selection of experimental validation candidates, we loosened the threshold for templates to *<* 0.5 to include more weak and medium binders. In the experiments with IEDB label-sanitized dataset in Section 2.3.2, we only included AVIB classifiers trained on the sanitized dataset in the ensemble classifier.
- **Clustering and representative selection:** The remaining samples were clustered by their 3-mer profiles using hierarchical clustering, with the number of clusters set to 75% of the size of the TCR sequence pool. For each cluster greater than 20, we chose the top 20 samples ranked by the ensemble binding score.
- **Ranking by interaction energy heuristics:** We calculated the Miyazawa-Jernigan energy heuristic using the pairwise energy matrix from the original 19 publication (Table VI) [34]. Specifically, for the TCR CDR3*β* and peptide pair, we used a sliding window of size 5 on each sequence and chose the lowest energy score among all possible interaction window pairs. Such heuristic is shown to distinguish between positive and negative TCRs for the MART-1 epitope (Fig. 3D). For experimental validation, we chose the TCR sequences with the lowest estimated energy among those with indistinguishable high binding scores (close to 1).

### 4.4 Computational structure and interaction energy prediction

#### 4.4.1 TCR-pMHC 3D structure prediction

We used the TCRmodel2 [43], a tailored pipeline specialized for TCR-pMHC structure prediction based on AlphaFold2 [63], to predict the TCR-pMHC complex structure for the wild-type reference TCR (start TCR2), three engineered variants (Eg2-1, Eg2-2, Eg2-3) and the two negative controls (Neg2-1, Neg2-2) with default parameters and using all five AlphaFold2 models.

#### 4.4.2 DSMBind fine-tuning and scoring

DSMBind [41] is an unsupervised scoring model of protein-protein interactions using denoising score matching, trained on general interaction complex structural data. Due to the distinct flat topology [64] and lower binding affinity [65] of TCR-pMHC interface compared to general protein-protein interactions, we fine-tuned DSMBind on curated TCR-pMHC structural data to capture specific principles governing TCR recognition, following the practice of several works dealing with TCR-pMHC structures [43, 66]. The fine-tuning data included known TCR-pMHC structures from the Protein Data Bank (PDB) and the predicted structures of their known binding mutants by Rosetta [42], provided by Shcherbinin et al. (2023) [67]. We used the structures after side chain repacking (“repacked”) and split the dataset into train/test/validation based on the templates, such that mutants derived from the same template belong to the same split. We treated the Rosetta-predicted energy score provided by the source dataset as the ground truth, and used the Spearman correlation between the DSMBind score and the ground truth energy score as the cross-validation metric during the fine-tuning. The fine-tuning was performed with learning rate 1.0 × 10^−4^ or 100 epochs. We observe improved correlation in the test set after fine-tuning, compared to the negative correlation from the pre-trained model (Supplementary).

The fine-tuned model was then applied to the predicted structures from 4.4.1. Specifically, we performed five TCRmodel2 runs for each system, each initialized with a distinct random seed (42, 123, 456, 789, 987) to account for the stochastic variability of the structural estimate of the flexible CDR loops, and selected the highest-ranking prediction in each run for downstream analysis.

#### 4.4.3 Binding free energy calculations from Rosetta

The top five TCRmodel2 outputs for each system were initially refined with the Rosetta FastRelax protocol with coordinate constraints. To enhance conformational sampling, additional rotameric states were explored during repacking, and 10,000 Monte Carlo trajectories were performed using the backrub algorithm [68], yielding 1,000 structures per system. Then, binding free energy calculations were carried out for the ensemble of structures using the Rosetta software suite [42]. Binding energies were computed with the InterfaceAnalyzer protocol, measuring the difference between the energy of the bound complex and the sum of the energies of the separated partners after side-chain repacking of both states. Final binding free energy values were obtained with the ddG mover by averaging results over all 1,000 complexes.

### 4.5 Molecular dynamics simulation

#### 4.5.1 System Building

Starting structures of HLA-A*02:01 bound to the ELAGIGILTV peptide and wild-type, engineered, or control TCRs were generated using TCRmodel2 [43]. For each system, the two highest-ranking TCRmodel2 predictions were selected for refinement, corresponding to models with confidence and i-pLDDT scores above 0.875. The selected models differed only in the docking orientation and incidence angle of TCR binding. Because TCRmodel2 outputs included a truncated HLA-A*02:01 structure (residues 1–180) and omitted the *β*2-microglobulin (*β*2m) chain, complete and structurally stable complexes were reconstructed by superimposing full-length HLA-A*02:01 and *β*2m from PDB ID 7RTD [69] onto the predicted models using PyMOL. The MHC and TCR were subsequently glycosylated at Asn86 and Asn405, respectively, in accordance with their known glycomic profiles [70]. A single complex-type N-glycan was modelled on the TCR, consistent with the predominant glycosignature observed in T cells [70].

#### 4.5.2 Molecular dynamics simulations

The built TCR-pMHC complexes, including *β*-microglobulin (*β*2m), MHC, and associated N-glycans, were subjected to molecular dynamics (MD) simulations. Protonation states of titratable residues were first determined at pH 7.4 using the PROPKA3 code [71]. All simulations were carried out using GROMACS 2022.5 [72], with the Amber ff14SB force field for proteins [73], GLYCAM06 for carbohydrates,[74] and the TIP3P water model [75]. Each system was placed in a cubic box with periodic boundary conditions, and Na^+^ and Cl^−^ ions were added to neutralize the overall charge and achieve a physiological salt concentration of 150 mM. Long-range electrostatic interactions were treated with the Particle Mesh Ewald (PME) method [76], while Lennard-Jones interactions were truncated at 10 °A. Energy minimization was performed for 30,000 steps, followed by heating to 298.15 K over 100 ps in the NVT ensemble. During this phase, a positional restraint of 5 kcal·mol^−1^·Å^−2^ was applied to all backbone atoms. Subsequently, equilibration was carried out in the NPT ensemble through six consecutive stages of 20 ps each, using progressively decreasing backbone restraints starting at 5, 4, 3, 2, and 1 kcal.mol^−1^·Å^−2^, followed by a final stage without restraints. The temperature was controlled using a Langevin thermostat with a collision frequency of 1.0 ps^−1^ and pressure maintained at 1 bar using a Parrinello-Rahman barostat [77, 78] with a relaxation time of 2 ps. Bond lengths involving hydrogen atoms were constrained using the LINCS algorithm [79]. For the production phase, three independent replicas were run for each model of each system, carrying out 20 ns per replica, resulting in a total of 120 ns of simulation per system.

#### 4.5.3 Molecular mechanics with generalized Born surface area

Binding free energies of the TCR–peptide:MHC complexes were estimated using MM/GBSA calculations following the approaches of Crean et al. (2022) [45] and Ribeiro-Filho et al. (2024) [80]. Calculations were performed with the gmx MMPBSA package [81] integrated into GROMACS 2022.5. For each replica, only the equilibrated portions of short trajectories were considered, corresponding to 2.5 ns per replica, with frames extracted every 10 ps. Prior to analysis, explicit water molecules and counterions were removed from all frames. The ionic strength was set to 0.15 M and the internal dielectric constant to 6.0, while all other parameters were kept at their default values. MM/GBSA calculations employed the GB-neck2 implicit solvent model [82]. Interfacial water molecules and entropy contributions were not included.

### 4.6 Cellular response experiments

#### 4.6.1 Generation of TCR reporter cell line (stable CD8-J2-TCR cells) and stimulation

TCR *β* chain cDNA, the nucleotide encoding self-cleaving P2A peptide, TCR *α* chain cDNA were assembled together in a linearized pMXs-IRES-puro retroviral vector (Cat. No. RTV-014, CELL BIOLABS INC) using the NEBuilder HiFi DNA Assembly Master Mix (New England Biolabs, E2621). The constructed plasmid vector, pMXs-TCR*β*-P2A-TCR*α*-IRES-puro, was utilized for retrovirus production. We used a retrovirus-packaging cell line (PhoenixGP-GaLV) [83] to produce a retrovirus for transducing TCR genes. The CD8-J2 cells retrovirally transduced with TCRs were cultured in the presence of puromycin (1 µg/ml) for selection (stable CD8-J2-TCR cells).

#### 4.6.2. Measuring the reactivity of the target TCR using transcriptionally active PCR (TAP) fragments

The TCR activity was assessed using a reporter cell line transiently expressing TCR through transcriptionally active PCR (TAP) fragments, as previously described [84]. Briefly, eBlocks genes (IDT) encoding variable regions of the TCR *α* and *β* chains were amplified by PCR. The amplified DNAs were assembled into linearized plasmid vectors containing a constant region of a TCR *α* or TCR *β* chain by the Gibson Assembly method. TAP fragments of TCR *α* and TCR *β* together with the pGL4.30 [luc2P/NFAT-RE/Hygro] vector (Promega E8481) were transfected into the ΔTCR*β* Jurkat cell-line J.RT3-T3.5 (ATCC Cat# TIB-153, RRID:CVCL 1316) expressing CD8*α* (CD8-J2)[85] by electroporation (Neon Electroporation System, Thermo Fisher) (CD8-J2-Luc-TCR). 5 × 10^4^ CD8-J2-Luc-TCR cells were cultured overnight in the presence of T2 cells (2 × 10^4^) with or without antigenic peptides. Activation of the NFAT reporter gene was measured by the Steady-Glo Luciferase Assay System (Promega).

#### 4.6.3 Data processing

Raw reporter assay readouts for each TCR were normalized by their respective mean anti-CD3 control signal, then averaged across three plates. At each peptide concentration, responses were compared with the template TCR (start TCR1 and start TCR2) using a one-sided t-test.

## References

[1] Bentley, G. A. & Mariuzza, R. A. The structure of the t cell antigen receptor. Annual review of immunology 14, 563–590 (1996).

[2] Sadelain, M., Riviére, I. & Riddell, S. Therapeutic t cell engineering. Nature 545, 423–431 (2017).

[3] Baulu, E., Gardet, C., Chuvin, N. & Depil, S. Tcr-engineered t cell therapy in solid tumors: State of the art and perspectives. Science advances 9, eadf3700 (2023).

[4] Hebeisen, M. et al. Identifying individual t cell receptors of optimal avidity for tumor antigens. Frontiers in immunology 6, 582 (2015).

[5] Aleksic, M. et al. Different affinity windows for virus and cancer-specific t-cell receptors: implications for therapeutic strategies. European journal of immunology 42, 3174–3179 (2012).

[6] Stone, J. D., Harris, D. T. & Kranz, D. M. Tcr affinity for p/mhc formed by tumor antigens that are self-proteins: impact on efficacy and toxicity. Current opinion in immunology 33, 16–22 (2015).

[7] Campillo-Davo, D., Flumens, D. & Lion, E. The quest for the best: how tcr affinity, avidity, and functional avidity affect tcr-engineered t-cell antitumor responses. Cells 9, 1720 (2020).

[8] Hoffmann, M. M. & Slansky, J. E. T-cell receptor affinity in the age of cancer immunotherapy. Molecular carcinogenesis 59, 862–870 (2020).

[9] Sprent, J. & Kishimoto, H. The thymus and central tolerance. Philosophical Transactions of the Royal Society of London. Series B: Biological Sciences 356, 609–616 (2001).

[10] Crean, R. M. et al. Molecular rules underpinning enhanced affinity binding of human t cell receptors engineered for immunotherapy. Molecular Therapy-Oncolytics 18, 443–456 (2020).

[11] Holler, P. D. et al. In vitro evolution of a t cell receptor with high affinity for peptide/mhc. Proceedings of the National Academy of Sciences 97, 5387–5392 (2000).

[12] Dunn, S. M. et al. Directed evolution of human t cell receptor cdr2 residues by phage display dramatically enhances affinity for cognate peptide-mhc without increasing apparent cross-reactivity. Protein Science 15, 710–721 (2006).

[13] Li, Y. et al. Directed evolution of human t-cell receptors with picomolar affinities by phage display. Nature biotechnology 23, 349–354 (2005).

[14] Schmitt, T. M., Stromnes, I. M., Chapuis, A. G. & Greenberg, P. D. New strategies in engineering t-cell receptor gene-modified t cells to more effectively target malignancies. Clinical cancer research 21, 5191–5197 (2015).

[15] Müller, T. R. et al. Targeted t cell receptor gene editing provides predictable t cell product function for immunotherapy. Cell Reports Medicine 2 (2021).

[16] Zhao, Y. et al. High-affinity tcrs generated by phage display provide cd4+ t cells with the ability to recognize and kill tumor cell lines. The Journal of Immunology 179, 5845–5854 (2007).

[17] Robbins, P. F. et al. Single and dual amino acid substitutions in tcr cdrs can enhance antigen-specific t cell functions. The Journal of Immunology 180, 6116– 6131 (2008).

[18] Croce, G. et al. Phage display enables machine learning discovery of cancer antigen–specific tcrs. Science advances 11, eads5589 (2025).

[19] Vazquez-Lombardi, R. et al. High-throughput t cell receptor engineering by functional screening identifies candidates with enhanced potency and specificity. Immunity 55, 1953–1966 (2022).

[20] Springer, I., Besser, H., Tickotsky-Moskovitz, N., Dvorkin, S. & Louzoun, Y. Prediction of specific tcr-peptide binding from large dictionaries of tcr-peptide pairs. Frontiers in immunology 11, 1803 (2020).

[21] Weber, A., Born, J. & Rodriguez Martínez, M. Titan: T-cell receptor specificity prediction with bimodal attention networks. Bioinformatics 37, i237–i244 (2021).

[22] Gao, Y. et al. Pan-peptide meta learning for t-cell receptor–antigen binding recognition. Nature Machine Intelligence 5, 236–249 (2023).

[23] Montemurro, A. et al. Nettcr-2.0 enables accurate prediction of tcr-peptide binding by using paired tcrα and β sequence data. Communications biology 4, 1060 (2021).

[24] Springer, I., Tickotsky, N. & Louzoun, Y. Contribution of t cell receptor alpha and beta cdr3, mhc typing, v and j genes to peptide binding prediction. Frontiers in immunology 12, 664514 (2021).

[25] Grazioli, F. et al. On tcr binding predictors failing to generalize to unseen peptides. Frontiers in immunology 13, 1014256 (2022).

[26] Watson, J. L. et al. De novo design of protein structure and function with rfdiffusion. Nature 620, 1089–1100 (2023).

[27] Madani, A. et al. Large language models generate functional protein sequences across diverse families. Nature biotechnology 41, 1099–1106 (2023).

[28] Liu, B. et al. Design of high-specificity binders for peptide–mhc-i complexes. Science 389, 386–391 (2025).

[29] Chen, Z. et al. T-cell receptor optimization with reinforcement learning and mutation polices for precision immunotherapy, 174–191 (Springer, 2023).

[30] Grazioli, F. et al. Attentive variational information bottleneck for tcr–peptide interaction prediction. Bioinformatics 39, btac820 (2023).

[31] Goncharov, M. et al. Vdjdb in the pandemic era: a compendium of t cell receptors specific for sars-cov-2. Nature methods 19, 1017–1019 (2022).

[32] Tickotsky, N., Sagiv, T., Prilusky, J., Shifrut, E. & Friedman, N. Mcpas-tcr: a manually curated catalogue of pathology-associated t cell receptor sequences. Bioinformatics 33, 2924–2929 (2017).

[33] Vita, R. et al. The immune epitope database (iedb): 2024 update. Nucleic Acids Research 53, D436–D443 (2025).

[34] Miyazawa, S. & Jernigan, R. L. Residue–residue potentials with a favorable contact pair term and an unfavorable high packing density term, for simulation and threading. Journal of molecular biology 256, 623–644 (1996).

[35] Elnaggar, A. et al. Prottrans: Toward understanding the language of life through self-supervised learning. IEEE transactions on pattern analysis and machine intelligence 44, 7112–7127 (2021).

[36] Chowell, D. et al. Tcr contact residue hydrophobicity is a hallmark of immunogenic cd8+ t cell epitopes. Proceedings of the National Academy of Sciences 112, E1754–E1762 (2015).

[37] Dhusia, K., Su, Z. & Wu, Y. A structural-based machine learning method to classify binding affinities between tcr and peptide-mhc complexes. Molecular immunology 139, 76–86 (2021).

[38] Wright, K. M. et al. Hydrophobic interactions dominate the recognition of a kras g12v neoantigen. Nature communications 14, 5063 (2023).

[39] Zhao, X. et al. Tuning t cell receptor sensitivity through catch bond engineering. Science 376, eabl5282 (2022).

[40] Trautmann, L. et al. Dominant tcr vα usage by virus and tumor-reactive t cells with wide affinity ranges for their specific antigens. European journal of immunology 32, 3181–3190 (2002).

[41] Jin, W. et al. Dsmbind: Se (3) denoising score matching for unsupervised binding energy prediction and nanobody design. bioRxiv 2023–12 (2023).

[42] Leaver-Fay, A. et al. in Rosetta3: an object-oriented software suite for the simulation and design of macromolecules, Vol. 487 545–574 (Elsevier, 2011).

[43] Yin, R. et al. Tcrmodel2: high-resolution modeling of t cell receptor recognition using deep learning. Nucleic Acids Research 51, W569–W576 (2023).

[44] Norden, D. M., Navia, C. T., Sullivan, J. T. & Doranz, B. J. The emergence of cell-based protein arrays to test for polyspecific off-target binding of antibody therapeutics, Vol. 16, 2393785 (Taylor & Francis, 2024).

[45] Crean, R. M., Pudney, C. R., Cole, D. K. & Van der Kamp, M. W. Reliable in silico ranking of engineered therapeutic tcr binding affinities with mmpb/gbsa. Journal of Chemical Information and Modeling 62, 577–590 (2022).

[46] Stadinski, B. D. et al. Effect of cdr3 sequences and distal v gene residues in regulating tcr–mhc contacts and ligand specificity. The Journal of Immunology 192, 6071–6082 (2014).

[47] Rudolph, M. G., Stanfield, R. L. & Wilson, I. A. How tcrs bind mhcs, peptides, and coreceptors. Annu. Rev. Immunol. 24, 419–466 (2006).

[48] Davis, M. M. & Bjorkman, P. J. T-cell antigen receptor genes and t-cell recognition. Nature 334, 395–402 (1988).

[49] Matsui, K., Boniface, J. J., Steffner, P., Reay, P. A. & Davis, M. M. Kinetics of t-cell receptor binding to peptide/i-ek complexes: correlation of the dissociation rate with t-cell responsiveness. Proceedings of the National Academy of Sciences 91, 12862–12866 (1994).

[50] Corse, E., Gottschalk, R. A., Krogsgaard, M. & Allison, J. P. Attenuated t cell responses to a high-potency ligand in vivo. PLoS biology 8, e1000481 (2010).

[51] Hebeisen, M. et al. Shp-1 phosphatase activity counteracts increased t cell receptor affinity. The Journal of clinical investigation 123, 1044–1056 (2013).

[52] Zhong, S. et al. T-cell receptor affinity and avidity defines antitumor response and autoimmunity in t-cell immunotherapy. Proceedings of the National Academy of Sciences 110, 6973–6978 (2013).

[53] Nijkamp, E., Ruffolo, J. A., Weinstein, E. N., Naik, N. & Madani, A. Progen2: exploring the boundaries of protein language models. Cell systems 14, 968–978 (2023).

[54] Hayes, T. et al. Simulating 500 million years of evolution with a language model. Science 387, 850–858 (2025).

[55] Graves, A. & Schmidhuber, J. Framewise phoneme classification with bidirectional lstm networks, Vol. 4, 2047–2052 (IEEE, 2005).

[56] Ren, J. et al. Likelihood ratios for out-of-distribution detection. Advances in neural information processing systems 32 (2019).

[57] Zong, B. et al. Deep autoencoding gaussian mixture model for unsupervised anomaly detection (2018).

[58] Li, T., Guo, H., Grazioli, F., Gerstein, M. & Min, M. R. Disentangled wasserstein autoencoder for t-cell receptor engineering. Advances in Neural Information Processing Systems 36, 73604–73632 (2023).

[59] Chen, S.-Y., Yue, T., Lei, Q. & Guo, A.-Y. Tcrdb: a comprehensive database for t-cell receptor sequences with powerful search function. Nucleic acids research 49, D468–D474 (2021).

[60] Schulman, J., Wolski, F., Dhariwal, P., Radford, A. & Klimov, O. Proximal policy optimization algorithms. arXiv preprint 1707.06347 (2017).

[61] Henikoff, S. & Henikoff, J. G. Amino acid substitution matrices from protein blocks. Proceedings of the National Academy of Sciences 89, 10915–10919 (1992).

[62] Lin, V. et al. Tcr3d 2.0: expanding the t cell receptor structure database with new structures, tools and interactions. Nucleic Acids Research 53, D604–D608 (2025).

[63] Jumper, J. et al. Highly accurate protein structure prediction with alphafold. nature 596, 583–589 (2021).

[64] Raybould, M. I., Nissley, D. A., Kumar, S. & Deane, C. M. Computationally profiling peptide: Mhc recognition by t-cell receptors and t-cell receptor-mimetic antibodies. Frontiers in Immunology 13, 1080596 (2023).

[65] Birnbaum, M. E. et al. Deconstructing the peptide-mhc specificity of t cell recognition. Cell 157, 1073–1087 (2014).

[66] Bradley, P. Structure-based prediction of t cell receptor: peptide-mhc interactions. elife 12, e82813 (2023).

[67] Shcherbinin, D. S., Karnaukhov, V. K., Zvyagin, I. V., Chudakov, D. M. & Shugay, M. Large-scale template-based structural modeling of t-cell receptors with known antigen specificity reveals complementarity features. Frontiers in Immunology 14, 1224969 (2023).

[68] Smith, C. A. & Kortemme, T. Backrub-like backbone simulation recapitulates natural protein conformational variability and improves mutant side-chain prediction. Journal of molecular biology 380, 742–756 (2008).

[69] Szeto, C. et al. Molecular basis of a dominant sars-cov-2 spike-derived epitope presented by hla-a* 02: 01 recognised by a public tcr. Cells 10, 2646 (2021).

[70] Vicente, M. M. et al. Mannosylated glycans impair normal t-cell development by reprogramming commitment and repertoire diversity. Cellular & Molecular Immunology 20, 955–968 (2023).

[71] Olsson, M. H., Søndergaard, C. R., Rostkowski, M. & Jensen, J. H. Propka3: consistent treatment of internal and surface residues in empirical p k a predictions. Journal of chemical theory and computation 7, 525–537 (2011).

[72] Abraham, M. J. et al. Gromacs: High performance molecular simulations through multi-level parallelism from laptops to supercomputers. SoftwareX 1, 19–25 (2015).

[73] Maier, J. A. et al. ff14sb: improving the accuracy of protein side chain and backbone parameters from ff99sb. Journal of chemical theory and computation 11, 3696–3713 (2015).

[74] Kirschner, K. N. et al. Glycam06: a generalizable biomolecular force field. carbohydrates. Journal of computational chemistry 29, 622–655 (2008).

[75] Jorgensen, W. L., Chandrasekhar, J., Madura, J. D., Impey, R. W. & Klein, M. L. Comparison of simple potential functions for simulating liquid water. The Journal of chemical physics 79, 926–935 (1983).

[76] Darden, T., York, D., Pedersen, L. et al. Particle mesh ewald: An n log (n) method for ewald sums in large systems. Journal of chemical physics 98, 10089–10089 (1993).

[77] Parrinello, M. & Rahman, A. Crystal structure and pair potentials: A moleculardynamics study. Physical review letters 45, 1196 (1980).

[78] Parrinello, M. & Rahman, A. Polymorphic transitions in single crystals: A new molecular dynamics method. Journal of Applied physics 52, 7182–7190 (1981).

[79] Hess, B., Bekker, H., Berendsen, H. J. & Fraaije, J. G. Lincs: A linear constraint solver for molecular simulations. Journal of computational chemistry 18, 1463– 1472 (1997).

[80] Ribeiro-Filho, H. V. et al. Exploring the potential of structure-based deep learning approaches for t cell receptor design. PLOS Computational Biology 20, e1012489 (2024).

[81] Valdés-Tresanco, M. S., Valdés-Tresanco, M. E., Valiente, P. A. & Moreno, E. gmx mmpbsa: a new tool to perform end-state free energy calculations with gromacs. Journal of chemical theory and computation 17, 6281–6291 (2021).

[82] Nguyen, H., Roe, D. R. & Simmerling, C. Improved generalized born solvent model parameters for protein simulations. Journal of chemical theory and computation 9, 2020–2034 (2013).

[83] Kondo, E. et al. Efficient generation of antigen-specific cytotoxic t cells using retrovirally transduced cd40-activated b cells. The Journal of Immunology 169, 2164–2171 (2002).

[84] Hamana, H., Shitaoka, K., Kishi, H., Ozawa, T. & Muraguchi, A. A novel, rapid and efficient method of cloning functional antigen-specific t-cell receptors from single human and mouse t-cells. Biochemical and biophysical research communications 474, 709–714 (2016).

[85] Ohta, R., Demachi-Okamura, A., Akatsuka, Y., Fujiwara, H. & Kuzushima, K. Improving tcr affinity on 293t cells. Journal of immunological methods 466, 1–8 (2019).

