## Supplementary materials for "Generative AI-Guided Design of High-Affinity T Cell Receptors" (from another version of this preprint: version 1, 2026-02-12)

#### 1 Model training details

##### 1.1 Fine-tuning of ProtBERT

We initialized the model with pre-trained ProtBERT [1] weights obtained from Hugging Face (https://huggingface.co/Rostlab/prot_bert) with all the default hyper-parameters. The model is trained on a combined corpus including the 7,552,869 CDR3$\beta$ sequences from both the TChard dataset [2] and TCRdb [3] for 3 epochs. The embedding of each sequence was obtained as the embedding of the first token.

##### 1.2 Fine-tuning of DSMBind

The domain-specific training data for TCR-pMHC structure consist of the Rosetta-predicted, relaxed and repacked structures by Shcherbinin et al. (2023) [4], derived from 28 TCR-pMHC structure templates from the Protein Data Bank (PDB) [5]. It contains 1585 models featuring 247 unique CDR$\alpha$ and 485 unique CDR$\beta$ sequences. We ablated key training strategy choices to evaluate their contributions to the ranking performance. Specifically, we compared two baselines: the original pre-trained model and the model trained from scratch on the Shcherbinin dataset, and several fine-tuned variants on either the dataset alone, or a mixture that includes the experimentally solved TCR-pMHC structures from the PDB to improve the reliability and realism.

To probe the robustness and potential data leakage, we evaluated under two splitting protocols: a random split or a stricter hard split by template. Model initialization used checkpoints pre-trained on either the protein-protein interaction dataset SKEMPI [6] or antibody structure dataset SAbDab [7], as provided by the author. During the unsupervised fine-tuning, the model saw only the input structures and remained agnostic to the ground-truth binding energy. For model selection, we used the Spearman correlation between the predicted scores and the ground truth on the validation set. Final comparisons were based on the alignment between the model ranking of the designed TCRs and the reporter cell assay readouts. Specifically, we evaluated five experimentally validated designs (three positives and two negative controls) and reported binding-score changes relative to the corresponding template (start_TCR2). Overall, fine-tuning on the SKEMPI-pretrained model with the hard-split cross-validation, especially when augmented with real PDB structures, yields the strongest agreement with the experimental rankings (Table S2 "Best ranking"). These results underscore the distinctive characteristics of the TCR-pMHC interfaces, and suggest a more stringent model selection criterion can improve the generalizability of protein-protein interaction prediction.

| **Hyperparameter** | **Value** |
|---|---|
| Threshold of $r_b$ (binding) | 0.9 |
| Threshold of $r_v$ (validity) | 1.2577 |
| Discount factor $\gamma$ | 0.9 |
| LSTM latent dimension | 256 |
| Hidden dimension of policy & value networks | 128 |
| Hidden layers of policy & value networks | 2 |
| Batch size | 256 |
| Clip range | 0.2 |
| Learning rate | $3 \times 10^{-4}$ |
| Value coefficient $\alpha_1$ | 0.5 |
| Entropy coefficient $\alpha_2$ | 0.01 |

**Table. S1** Shared hyperparameters across `TCRPPO2` variants.

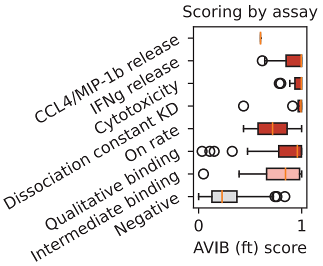


**Fig. S1** The prediction results of the fine-tuned AVIB classifier (unsanitized) on IEDB test samples by experimentally validated annotation labels.

| Model | Eg2-1 | Eg2-2 | Eg2-3** | Neg2-1 | Neg2-2 |
|---|---|---|---|---|---|
| Pretrained (SKEMPI) | -55.82 | -18.42 | -28.89 | -7.66 | -36.21 |
| Pretrained (SAbDAb) | | | | | |
| *From scratch* | | | | | |
| TCR | 15.02 | 22.20 | 2.99 | 22.69 | 7.74 |
| TCR+PDB | -66.05 | -27.18 | -69.52 | -34.39 | 1.87 |
| TCR (hard-split) | -78.40 | -30.78 | -81.04 | -41.43 | -2.36 |
| *Fine-tuned* | | | | | |
| TCR | -305.06 | -179.97 | -395.39 | -5.70 | 188.05 |
| TCR+PDB | -114.33 | -81.03 | -171.76 | 26.77 | 100.20 |
| TCR (hard-split) | -158.47 | -96.71 | -203.27 | 2.86 | 95.63 |
| Best ranking | 2 | 3 | 1 | 4 | 5 |
| Average ranking | 2.17 | 3.50 | 1.00 | 3.83 | 4.50 |
| Experimental ranking | 3 | 2 | 1 | 4.5 | 4.5 |

**Table. S2** DSMBind ranking of five experimentally validated TCR designs under different training strategies, reported as predicted binding-score changes relative to the corresponding template. Rankings with the highest agreement with the experimental reporter cell assay are underlined.

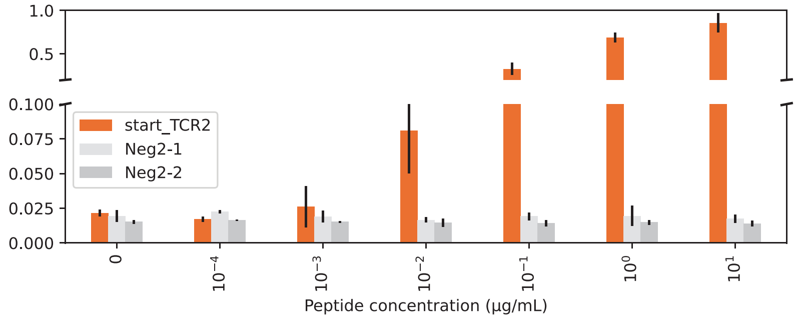


**Fig. S2** The reporter cell assay results for the selected initial (start_TCR2) and selected negative TCRs (Neg2-1, Neg2-2).

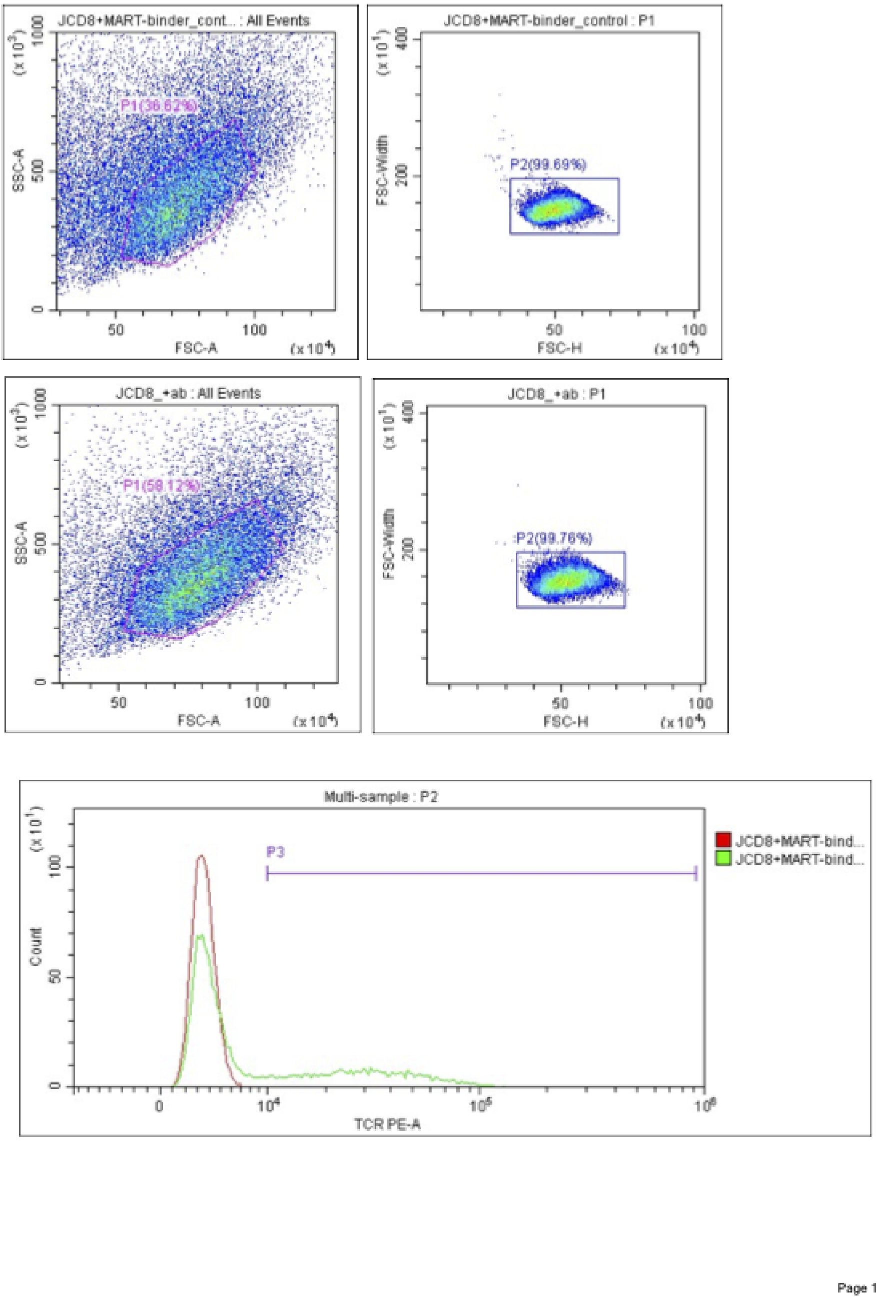


**Fig. S3** Flow cytometry validation of start_TCR1 (MART1 binder) TCR expression in Jurkat cells following the transfection.

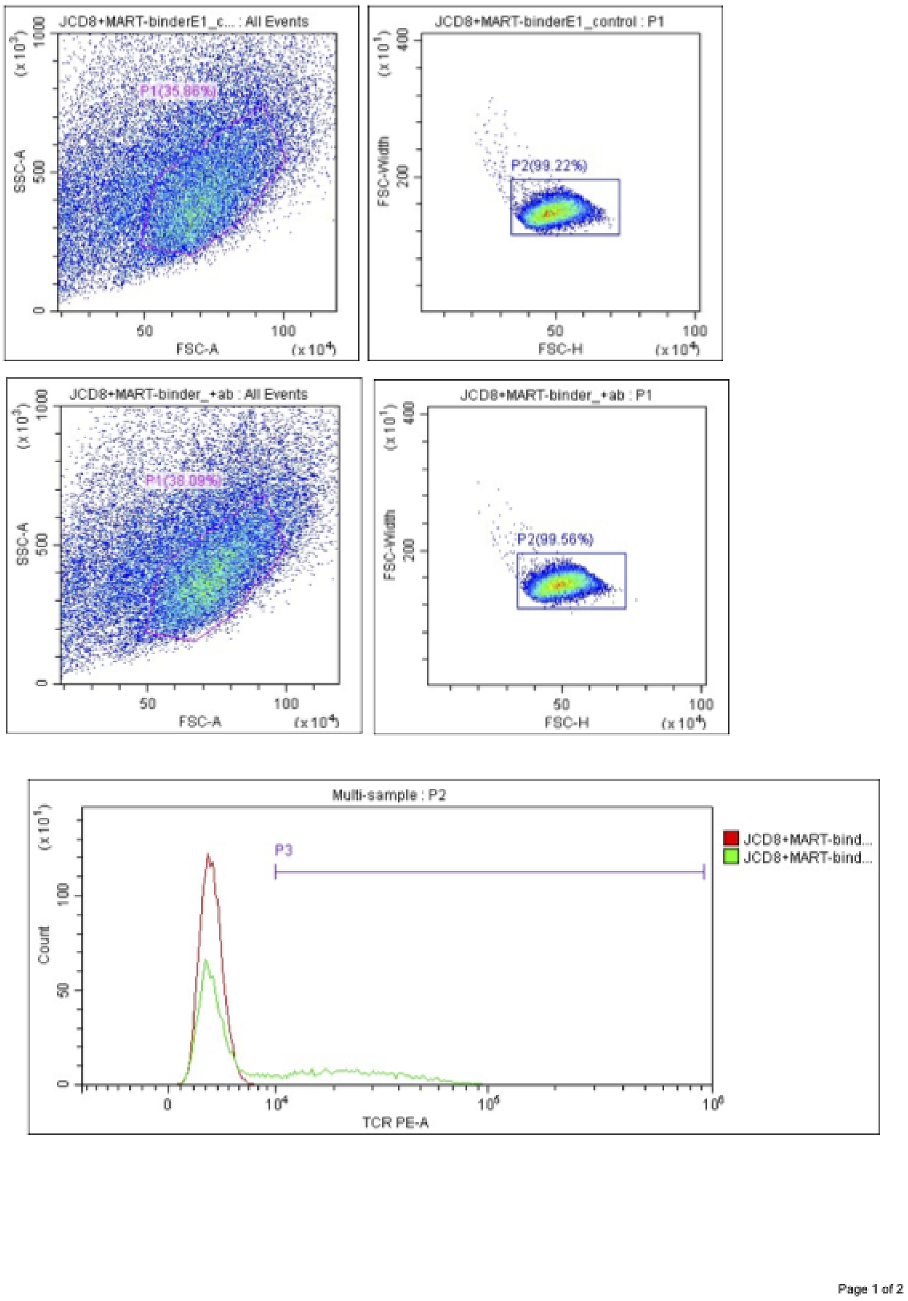


**Fig. S4** Flow cytometry validation of Eg1-1 (MART1 binder E1) TCR expression in Jurkat cells following the transfection.

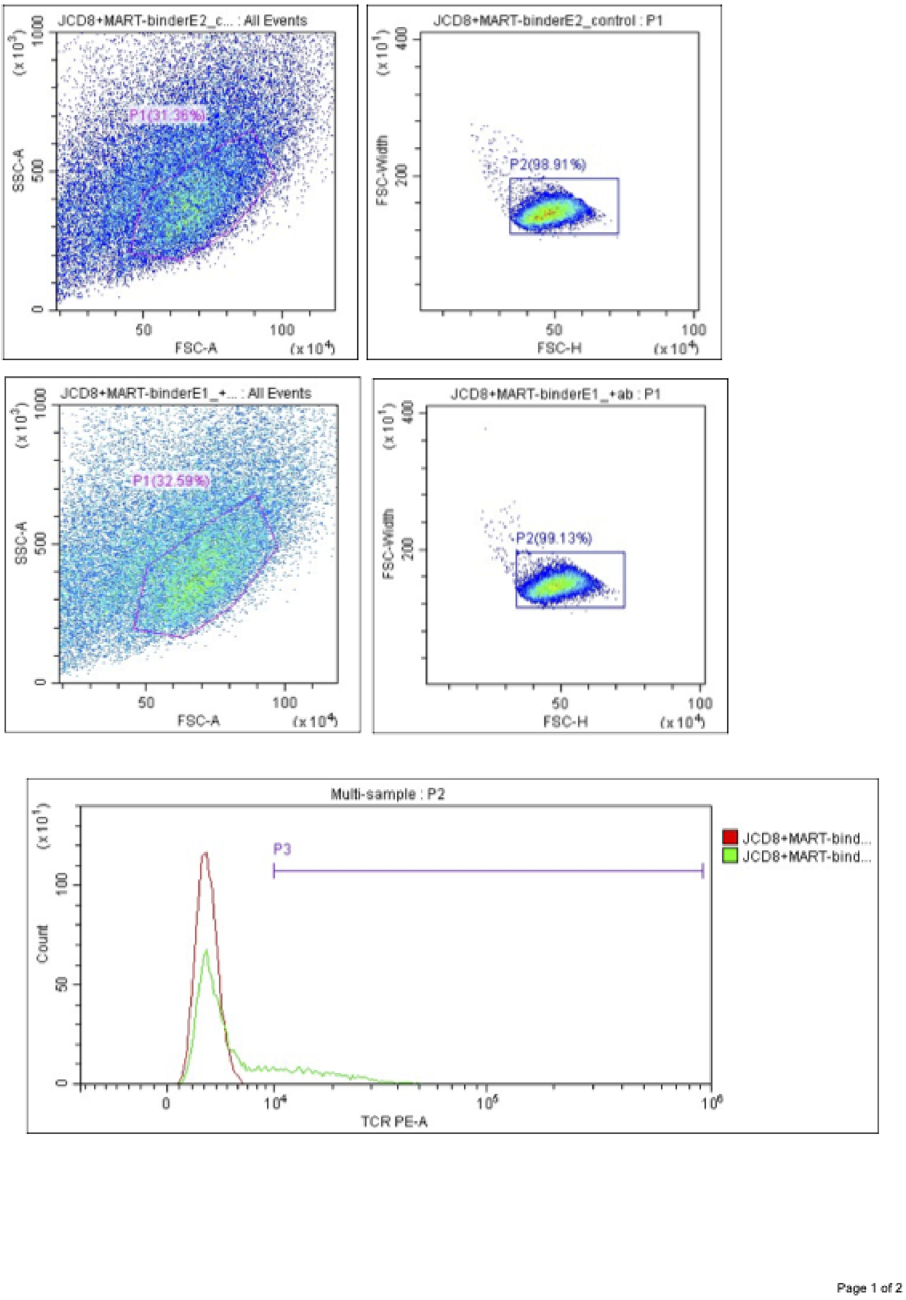


**Fig. S5** Flow cytometry validation of Eg1-2 (MART1 binder E2) TCR expression in Jurkat cells following the transfection.

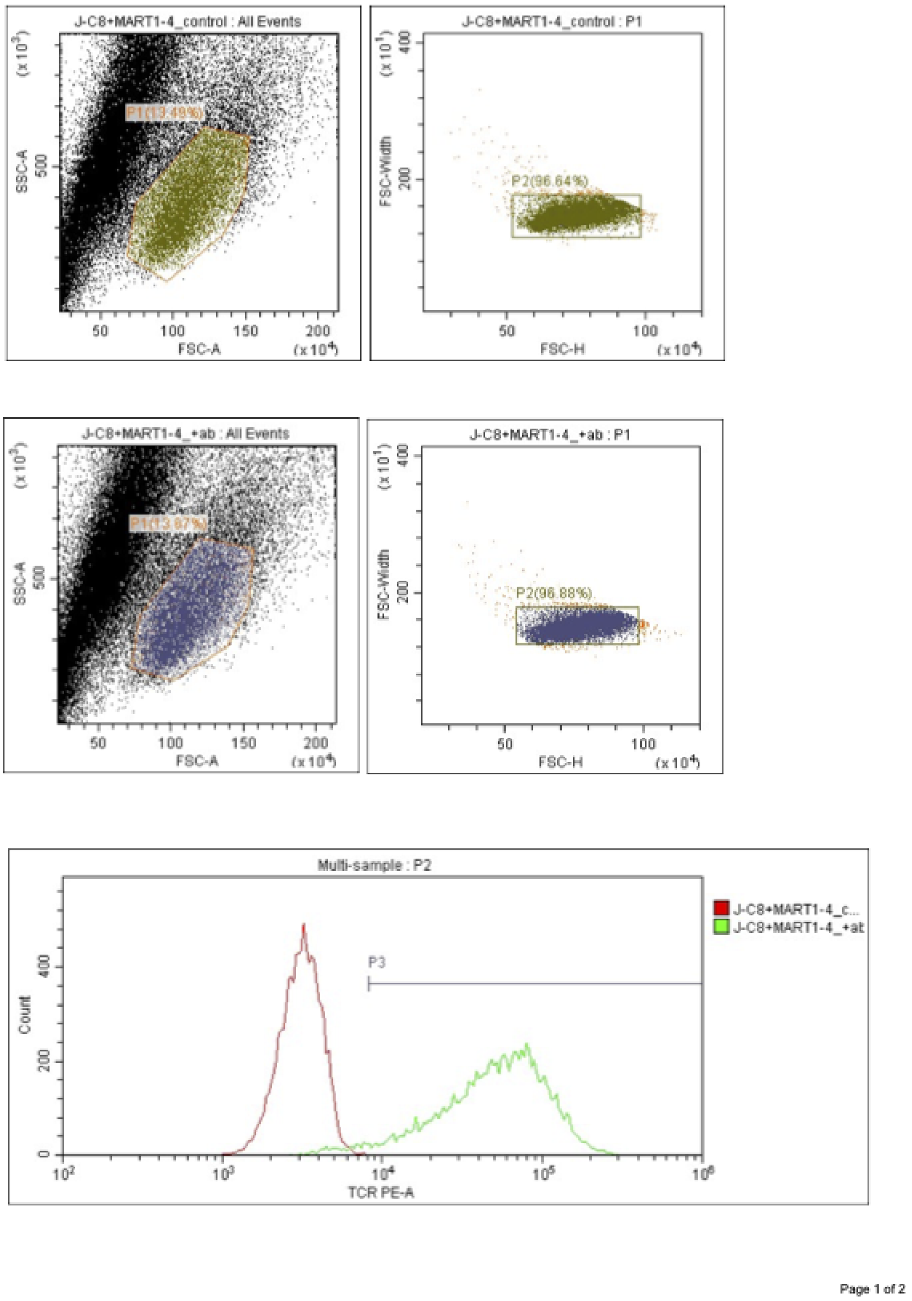


**Fig. S6** Flow cytometry validation of start_TCR2 (MART1-4) TCR expression in Jurkat cells following the transfection.

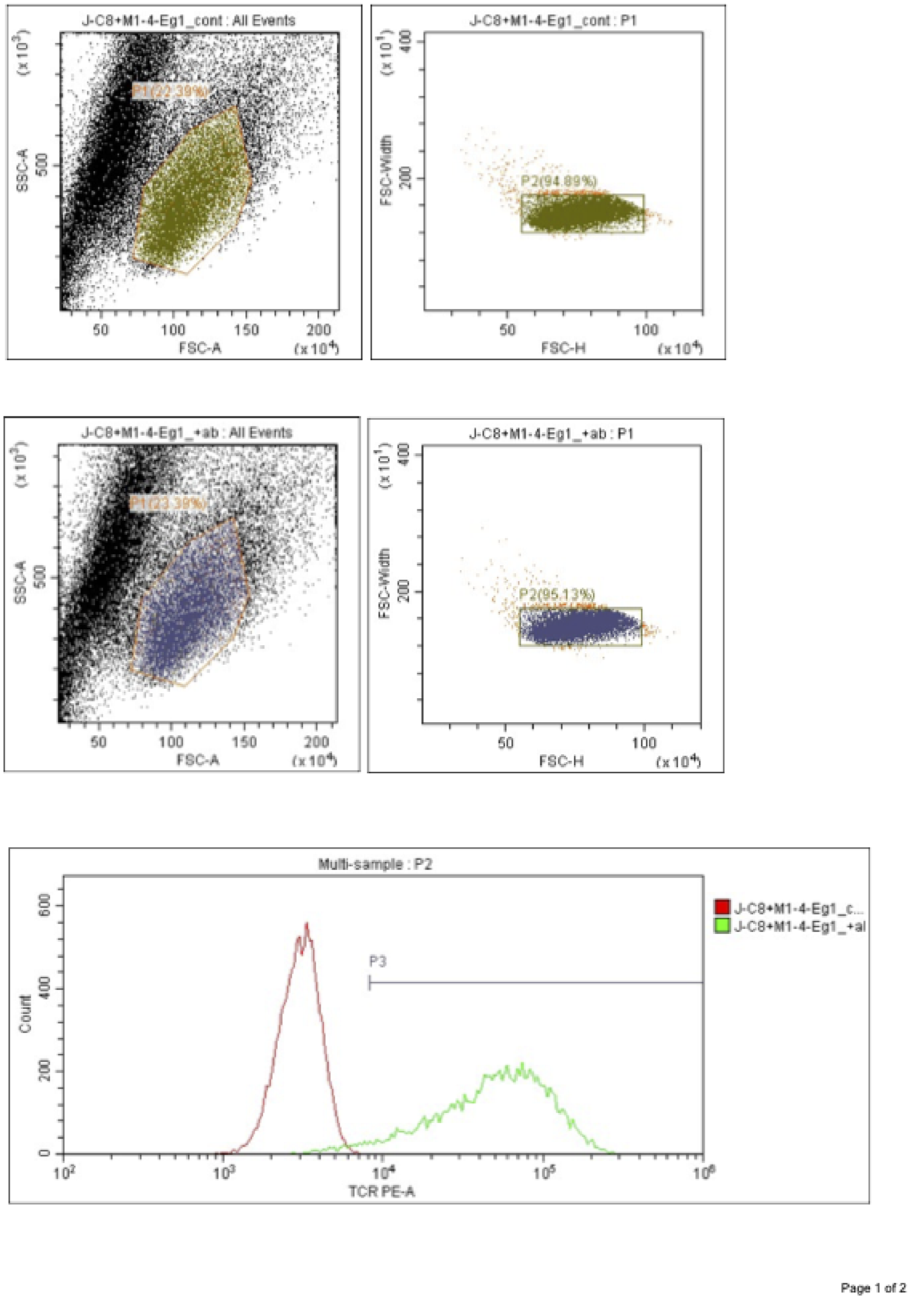


**Fig. S7** Flow cytometry validation of Eg2-1 (MART1-4-Eg1) TCR expression in Jurkat cells following the transfection.

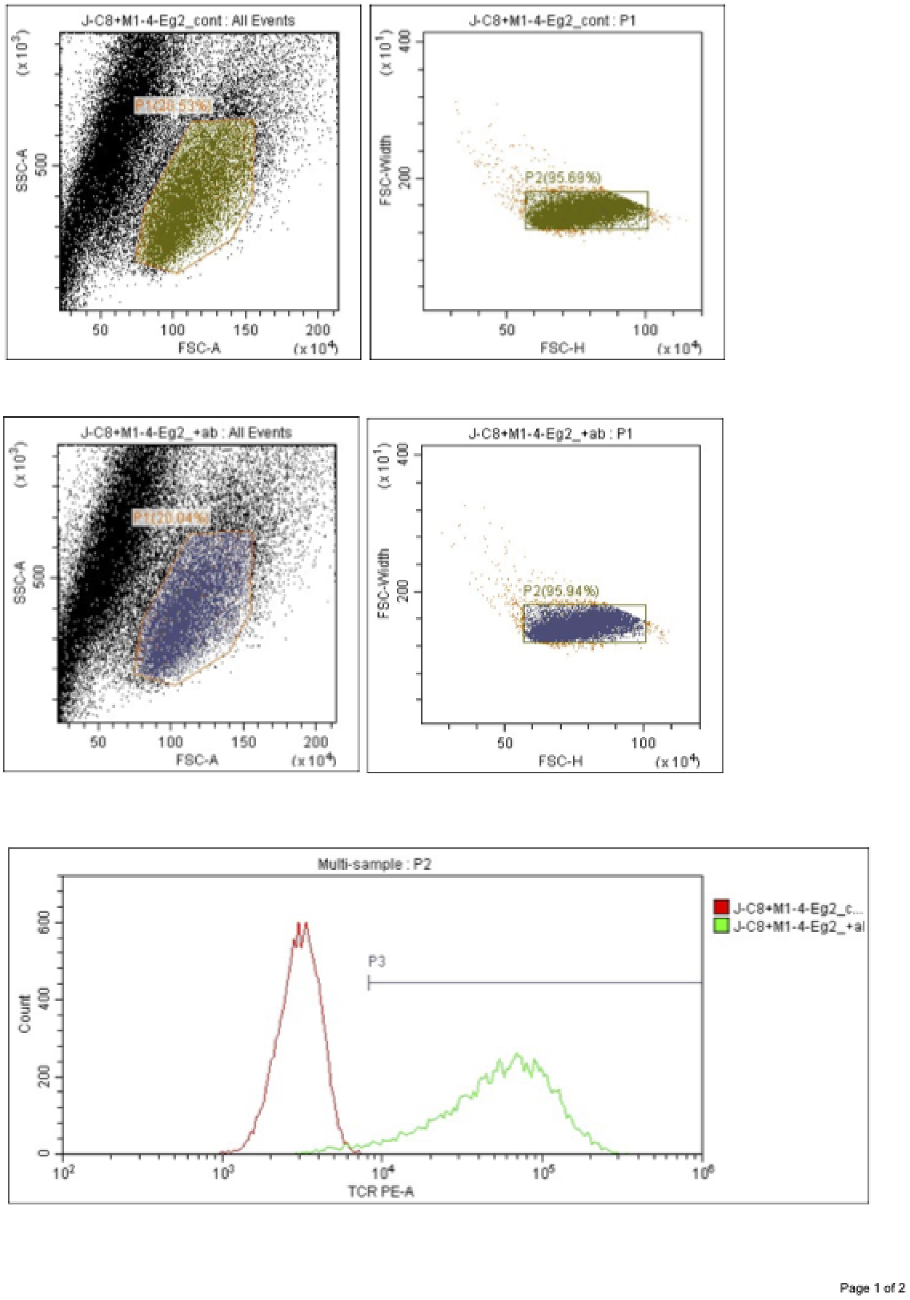


**Fig. S8** Flow cytometry validation of Eg2-2 (MART1-4-Eg2) TCR expression in Jurkat cells following the transfection.

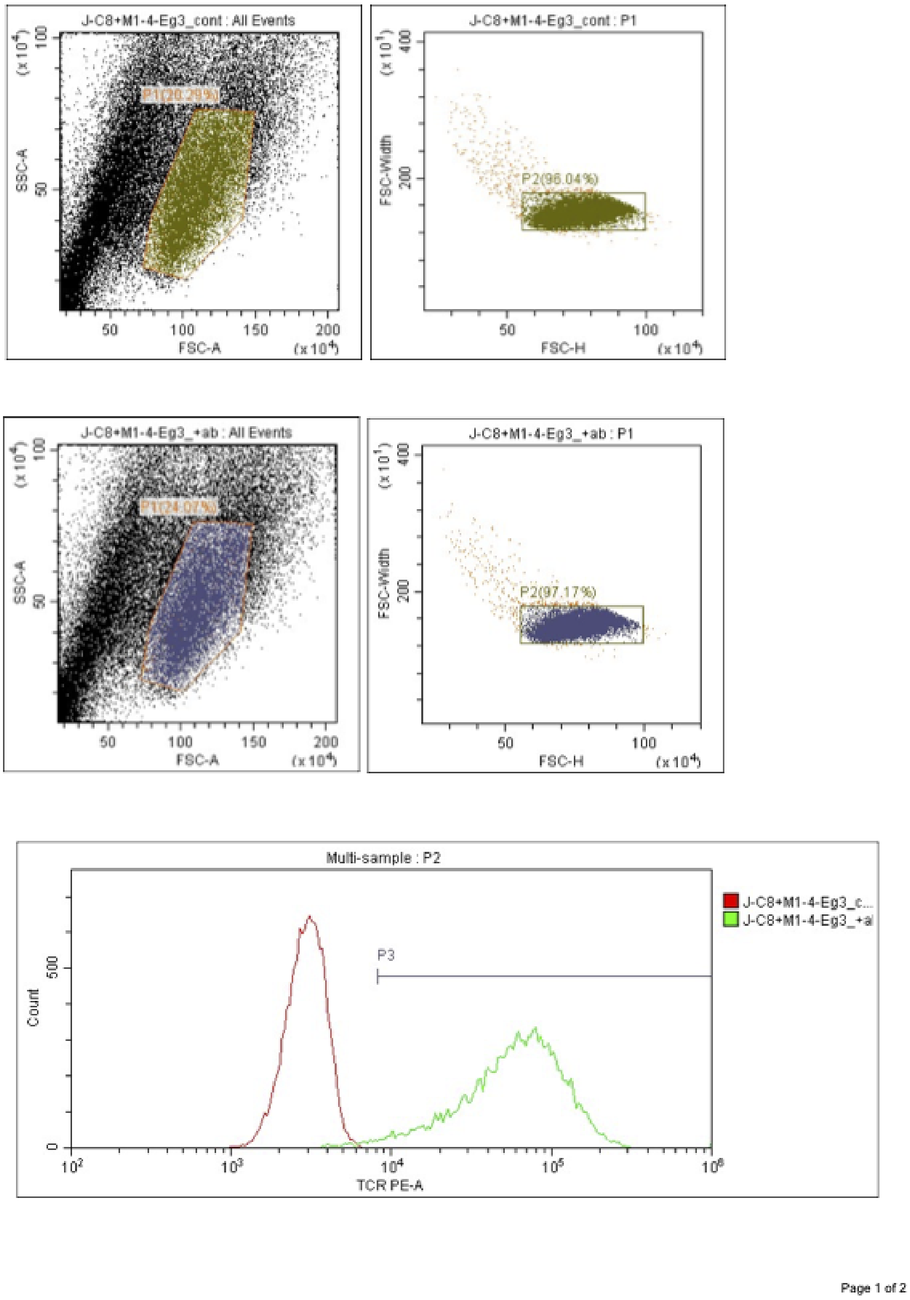


**Fig. S9** Flow cytometry validation of Eg2-3 (MART1-4-Eg3) TCR expression in Jurkat cells following the transfection.

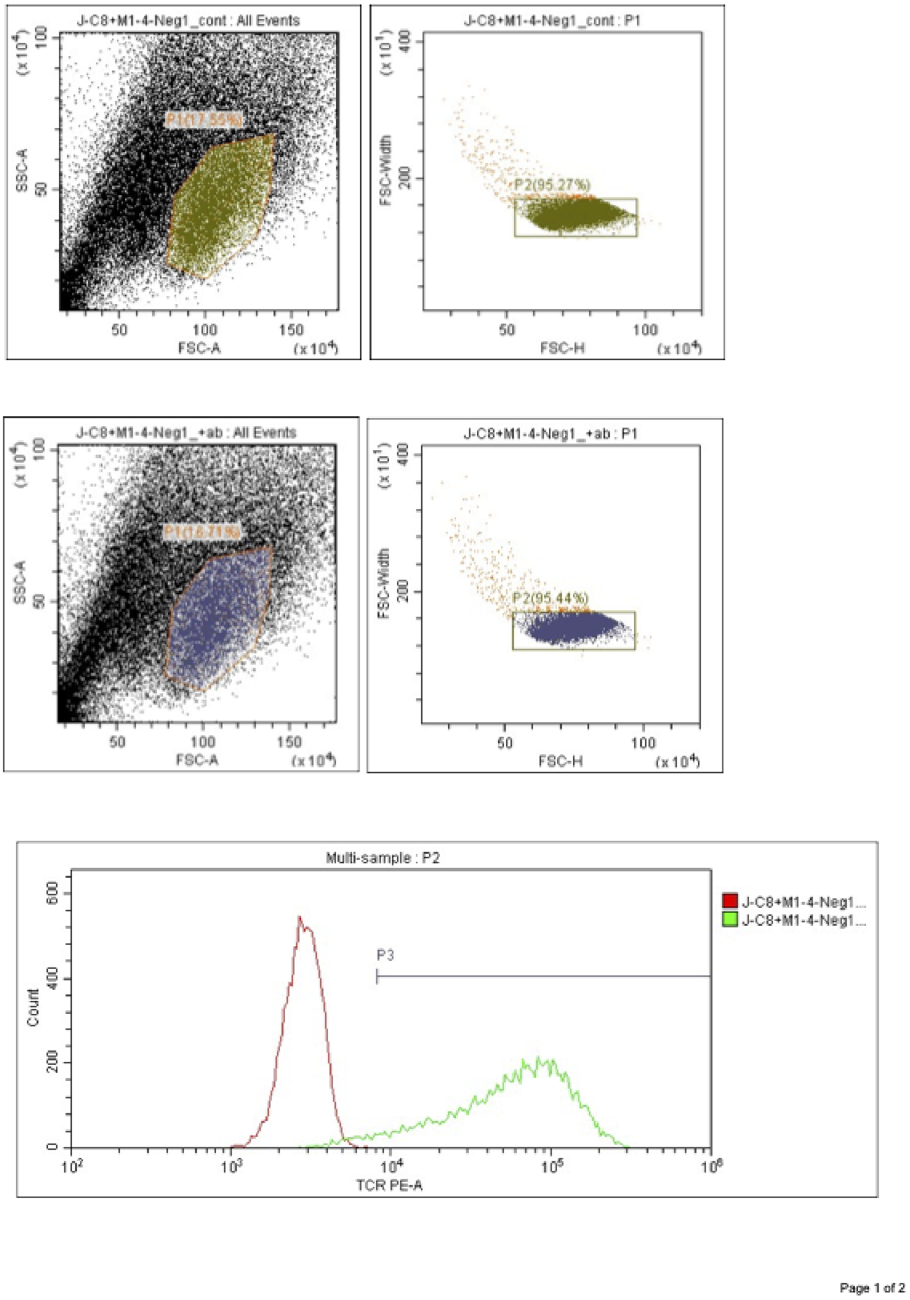


**Fig. S10** Flow cytometry validation of Neg2-1 (MART1-4-Neg1) TCR expression in Jurkat cells following the transfection.

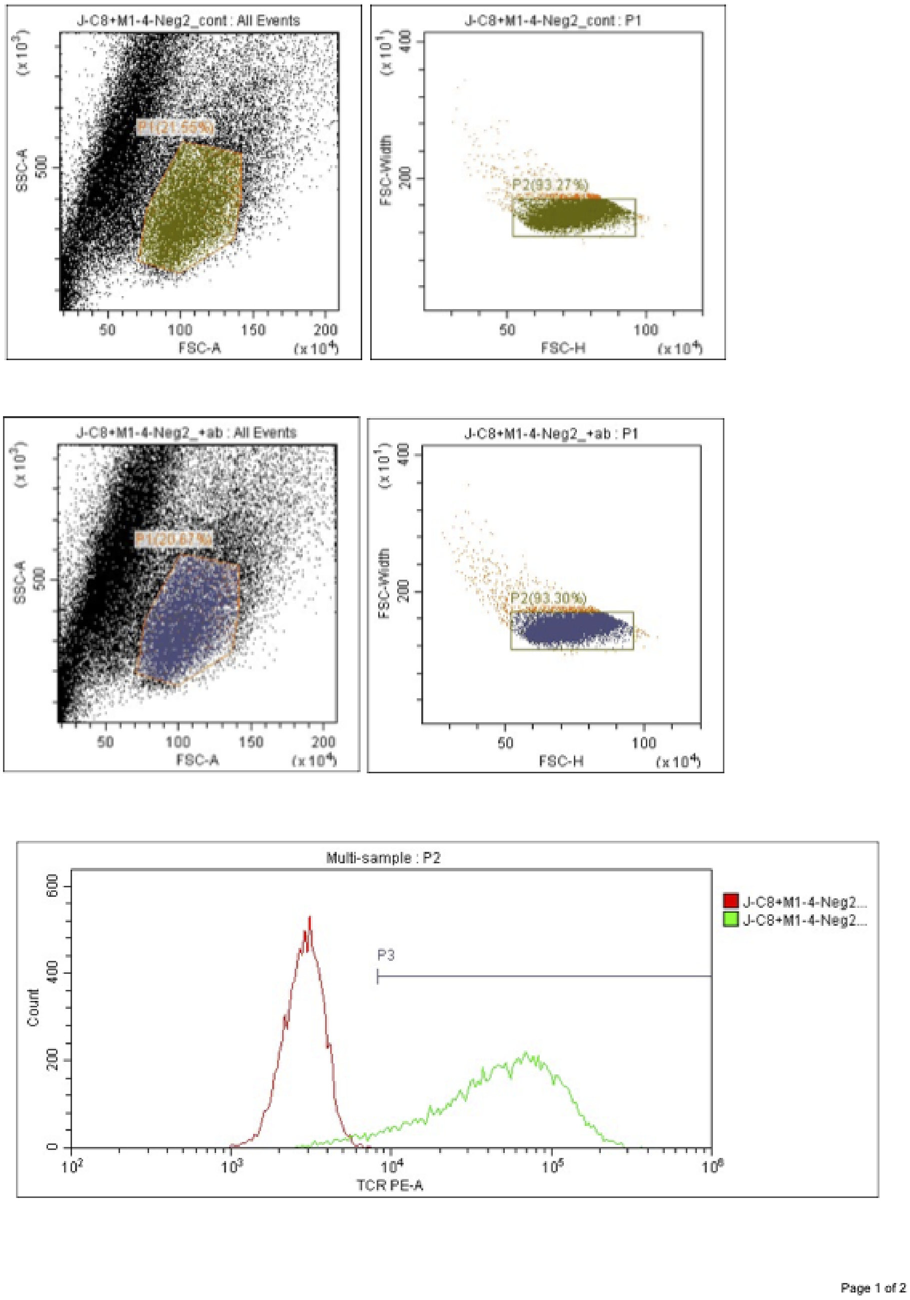


**Fig. S11** Flow cytometry validation of Neg2-2 (MART1-4-Neg2) TCR expression in Jurkat cells following the transfection.

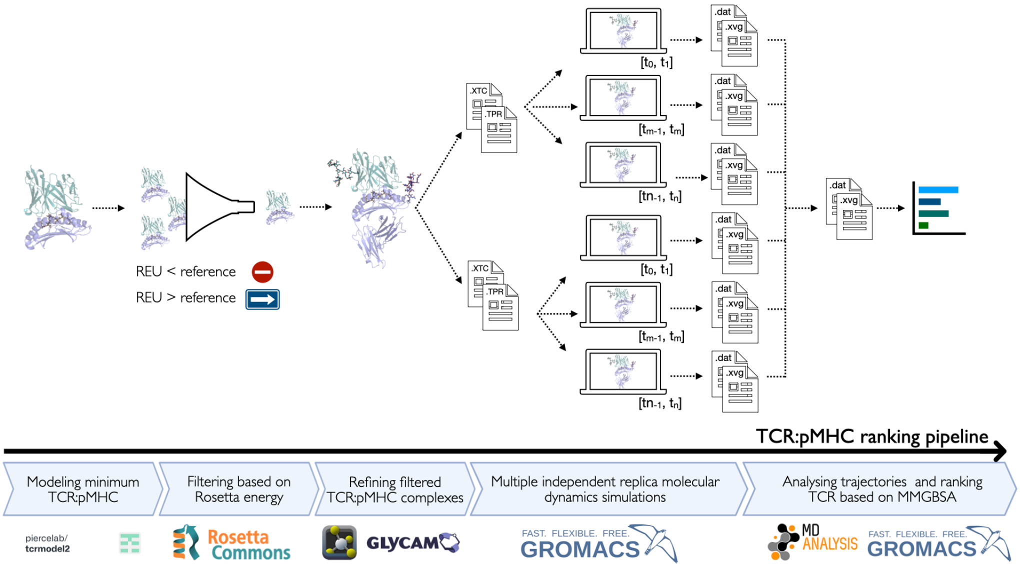


**Fig. S12 Computational pipeline for TCR:pMHC ranking.** Structural models of TCR:pMHC complexes were generated using TCRmodel2 and filtered based on Rosetta energy (REU). Complexes with favorable energy scores were refined using Rosetta Commons and GLYCAM, then subjected to multiple independent replica molecular dynamics simulations using GROMACS. Trajectories were analyzed with MDAnalysis and GROMACS to compute MM-GBSA binding energies for TCR ranking.

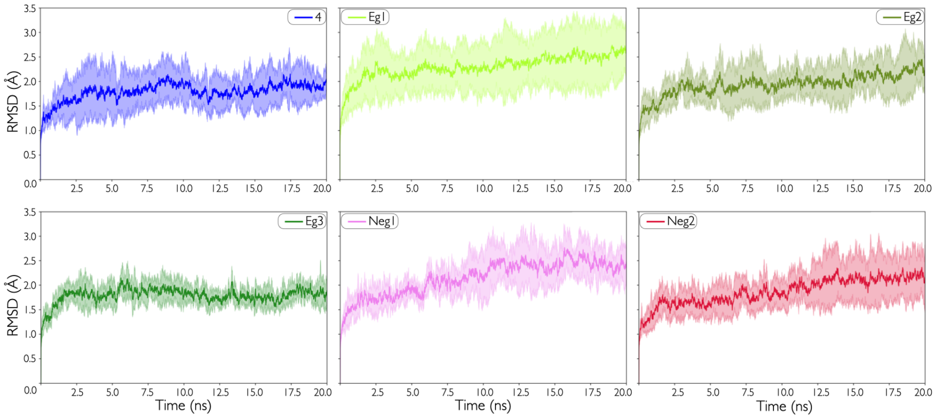


**Fig. S13 Root mean square deviation (RMSD) of TCR:pMHC complexes during molecular dynamics simulations** Backbone RMSD as a function of simulation time for systems 4, Eg1, Eg2, Eg3 (positive binders) and Neg1, Neg2 (negative controls). Solid lines represent the mean across replicas; shaded areas indicate standard deviation. All systems equilibrate within the first 2.5 ns and remain stable throughout the 20 ns production runs.

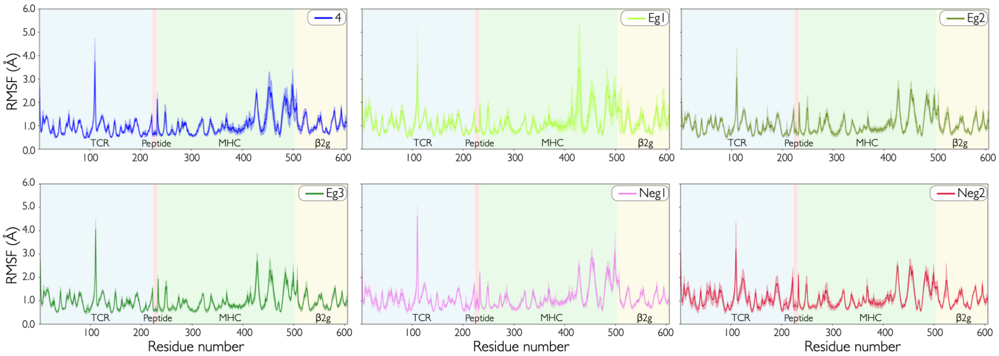


**Fig. S14 Per-residue root mean square fluctuation (RMSF) profiles of TCR:pMHC complexes.** Root mean square fluctuation per residue averaged across replicas. Regions corresponding to TCR (blue background), peptide (pink), MHC (yellow), and $\beta$2-microglobulin (green) are indicated. Higher flexibility is observed in TCR CDR loops and terminal regions across all systems.

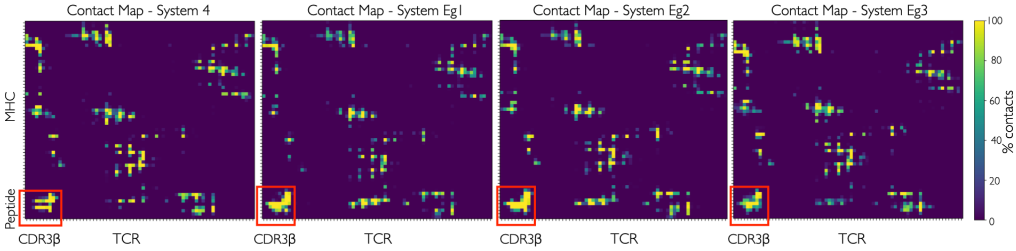


**Fig. S15 Contact frequency maps between TCR and pMHC.** Percentage of simulation frames in which residue pairs maintain contact (¡4.5 Å) for systems 4, Eg1, Eg2, and Eg3. The red box highlights CDR3$\beta$–peptide interactions, which are consistently maintained across all positive binders, supporting stable TCR engagement with the peptide epitope.

### References


[1] Elnaggar, A. *et al.* Prottrans: Toward understanding the language of life through self-supervised learning. *IEEE transactions on pattern analysis and machine intelligence* **44**, 7112–7127 (2021).

[2] Grazioli, F. *et al.* On tcr binding predictors failing to generalize to unseen peptides. *Frontiers in immunology* **13**, 1014256 (2022).

[3] Chen, S.-Y., Yue, T., Lei, Q. & Guo, A.-Y. Tcrdb: a comprehensive database for t-cell receptor sequences with powerful search function. *Nucleic acids research* **49**, D468–D474 (2021).

[4] Shcherbinin, D. S., Karnaukhov, V. K., Zvyagin, I. V., Chudakov, D. M. & Shugay, M. Large-scale template-based structural modeling of t-cell receptors with known antigen specificity reveals complementarity features. *Frontiers in Immunology* **14**, 1224969 (2023).

[5] Berman, H. M. *et al.* The protein data bank. *Nucleic acids research* **28**, 235–242 (2000).

[6] Jankauskaitė, J., Jiménez-García, B., Dapkūnas, J., Fernández-Recio, J. & Moal, I. H. Skempi 2.0: an updated benchmark of changes in protein–protein binding energy, kinetics and thermodynamics upon mutation. *Bioinformatics* **35**, 462–469 (2019).

[7] Schneider, C., Raybould, M. I. & Deane, C. M. Sabdab in the age of biotherapeutics: updates including sabdab-nano, the nanobody structure tracker. *Nucleic acids research* **50**, D1368–D1372 (2022).
